# Autonomous biogenesis of the entire protein translation machinery excluding ribosomes

**DOI:** 10.1101/2024.10.20.619270

**Authors:** Matthaeus Schwarz-Schilling, Aurore Dupin, Noa Avidan, Yoav Barak, Yoshihiro Shimizu, Shirley S. Daube, Roy H. Bar-Ziv

## Abstract

Recreating the conditions for autonomous biogenesis of the protein translation machinery is fundamental to our understanding of living systems and is essential for the assembly of minimal cell models. All components of the translation machinery, including the ribosomes, translation factors and aminoacyl tRNA synthetases, are made of proteins and are therefore involved in their own synthesis, posing a unique challenge for self-biogenesis. Here, we created physicochemical conditions for autonomous biogenesis of all the translation machinery, excluding the ribosome. We surface-immobilized synthetic genes coding for all thirty components as densely packed DNA brushes forming a reaction center that localizes, concentrates and catalyzes their simultaneous synthesis. To demonstrate their activity, we first determined empirically the minimal critical concentration of each translation protein required to initiate its own self-biogenesis in bulk solution. We then assembled a minimal gene expression reaction with all translation proteins at their critical concentrations. Under these dilute conditions, reporter proteins were not synthesized unless the DNA brushes encoded all the translation proteins, thereby demonstrating their co-synthesis, functionality and engagement in their own synthesis. This scenario of a complex biochemical process that amplifies itself can be generalized and extended to impact our understanding toward the design of autonomous self-replicating biological system.

## Introduction

All proteins in all kingdoms of life are synthesized by a universal protein translation machinery consisting of the ribosome and some thirty auxiliary proteins, which facilitate efficient cycles of protein synthesis. The protein translation machinery orchestrates the cellular protein repertoire by controlling the synthesis rate of every protein, including their own, in response to genetic and epigenetic signals, environmental cues, and against protein degradation^1^. As such, and given a supply of nutrients, the translation machinery can produce and regulate its own components and is, therefore, a central and essential part in any realization of a minimal model of a cell that can autonomously propagate, evolve, and maintain homeostasis based on a genetic program^2^. Two more cellular processes, DNA replication and RNA transcription, are secondary in significance for the reconstitution of an autonomous propagating cell model, as they are driven by protein enzymes, thereby themselves dependent on the biogenesis of the translation machinery.

The protein translation machinery has been reconstituted outside a cell just over two decades ago, using a minimal set of purified proteins at biochemical conditions that support multiple translation cycles^3^ (Fig. 1a, b). Termed the PURE system, for protein synthesis using recombinant elements, this cell-free bacterial platform has been shown throughout the years to support the synthesis of a myriad of proteins^4–7^. Finding conditions for the PURE system to amplify its own protein components in a functional form would mark an essential milestone toward an autonomous protein-based artificial cell^8,9^ (Fig. 1a, b). This entails the simultaneous synthesis of dozens of protein building blocks of the ribosomes and all the auxiliary protein machinery within the minimal and dilute conditions of the PURE system, unfavorable for multicomponent assembly. While improved versions of the PURE system with increased protein yields have been established^10–13^, such enriched conditions would challenge the detection of functional nascent components within the saturating background of the purified proteins that created them. Indeed, self-synthesis and functionality of PURE components has been demonstrated so far for only a subset of the auxiliary translation proteins^14–17^.

**Figure 1.**
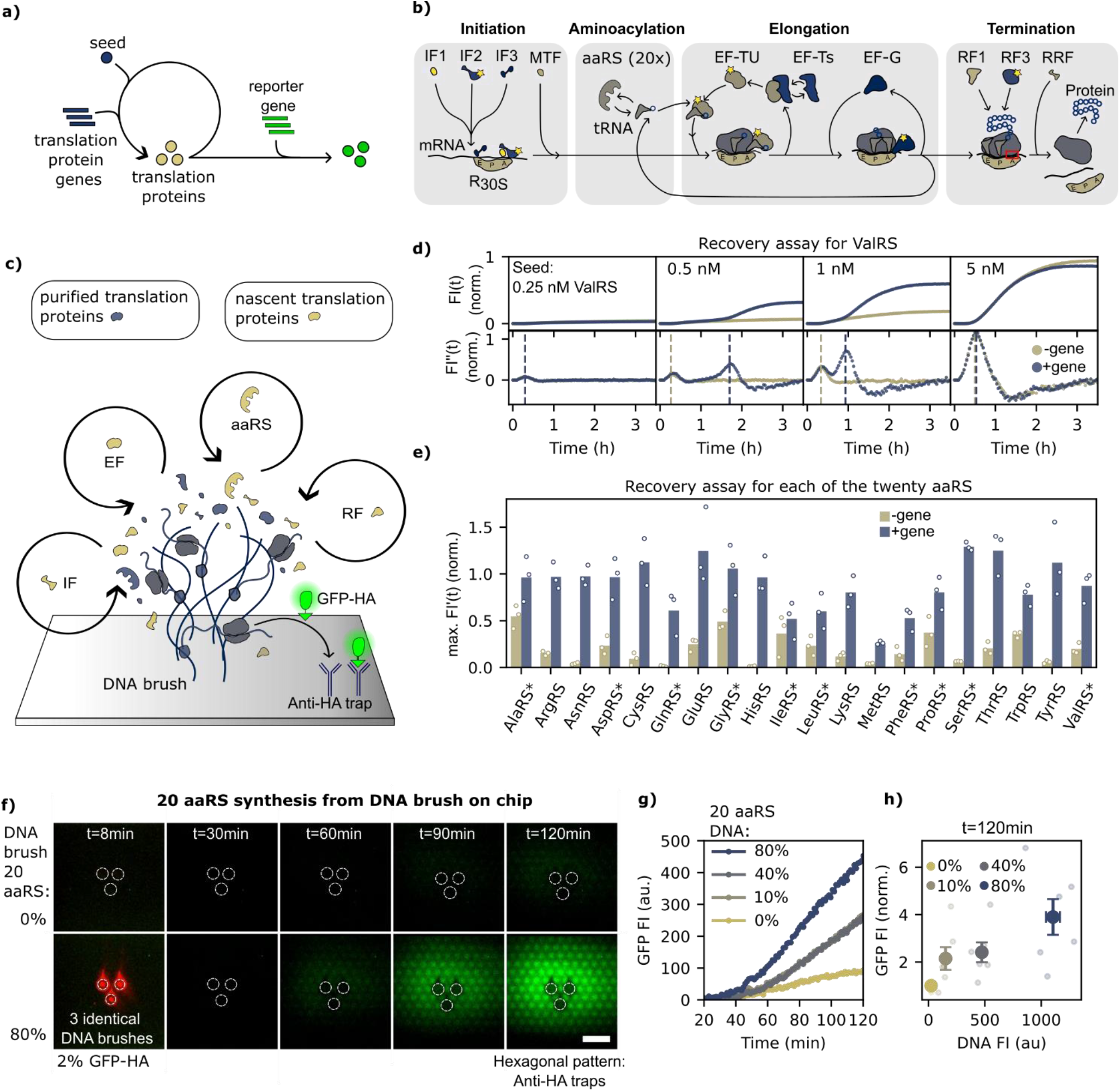
Biogenesis of twenty aaRSs from DNA brushes. **a)** Scheme: recovery of gene expression by self-synthesis of the translation protein from genes. **b)** Scheme: protein synthesis in the PURE system showing the 30 proteins involved in translation: initiation factors (IF), aminoacyl-tRNA synthetases (aaRS), elongation factors (EF), and release and recycling factors (RF). **c)** Scheme: DNA brush immobilized on a surface. Localized expression machinery (dark blue) starts the expression of nascent translation proteins (yellow), which concomitantly accumulate around the brush and promote self-synthesis. **d-e)** PURE recovery measurements in a microplate well. **d)** Fluorescence intensity time traces, FI(t), from a ΔValRS-PURE with different amounts of seeds +/−ValRS DNA (top) and the 2^nd^ derivative of their FI(t) (bottom). Fluorescence intensity (FI) are normalized to the max. Fl (top) or max. FI′′ (t) (bottom) of a full PURE. Dashed lines at t = max. FI′′ (t). [DNA_GFP_] = 1 nM, [DNA_ValRS_] = 1.5 nM. **e)** Bar plot: average normalized max. rate of ΔP_i_-PURE for 20 aaRS +/−P_i_-DNA. The ten aaRSs, which require an external seed to kick-start the self-synthesis, are marked with an asterisk (*). See Table S1. **f-h)** Biogenesis of all 20 aaRS from a DNA brush on a silica chip. **f)** TIRF microscopy images of a hexagon-patterned surface with traps for a GFP reporter (green channel) and three-DNA brush clusters (white dotted circles). Top row: DNA brushes code for a non-interacting control protein (DNA not labeled). Bottom row: DNA brushes code for twenty aaRSs (DNA-red labeled, red-channel TIRF signal overlayed in the left image). All brushes contain 2% GFP-HA genes. Upon exposure to a ΔPURE seeded according to the recovery assay in e) S1), surface-bound GFP-HA accumulated around DNA brushes depending on the fraction of aaRS genes. Scale bar = 200 μm. **g)** Average GFP Fl(t) from TIRF microscopy images for DNA brushes with different fractions of aaRS genes. **h)** Normalized average GFP Fl and its s.e.m. at t=120 min. from TIRF microscopy as a function of the DNA Fl (aaRS + GFP DNA) for different DNA brushes. n=6. Values are normalized to the average GFP signal of a DNA brush with 0% aaRS genes.

Recently, we presented a cell-free scenario for localization and confinement of gene expression reactions at surface-immobilized DNA brushes, facilitating co-synthesis and assembly of twenty-seven parts of the E. coli small ribosomal subunit (SSU)^18^. The high-density gene brushes localize the PURE gene expression machinery, with a concomitant accumulation of nascent RNA and protein products, increasing their local concentration and facilitating macromolecular interactions^19^. As the many bottlenecks towards reconstitution of the biogenesis of the large ribosomal subunit (LSU) have yet to be overcome, we focus here on the rest of the translation machinery, including twenty aminoacyl transfer RNA (tRNA) synthetases (aaRS) and ten translation factors (Fig. 1b), and demonstrate, using the DNA brush methodology, the simultaneous self-biogenesis of all thirty proteins and their active involvement in gene expression.

While ribosomal parts form a tight complex that could be trapped on the surface to indicate assembly^18^, the aaRSs and translation factors are only loosely associated with the ribosome, requiring a functional assay to indicate their self-synthesis and functionality in a process far from equilibrium. We immobilized all thirty genes coding for the translation proteins as dense DNA brushes and detected with high sensitivity by Total Internal Fluorescence (TIRF) microscopy the fluorescence of a reporter Green Fluorescent Protein (GFP) trapped on the surface (Fig. 1c). The autonomous localization of the PURE machinery to the high DNA density allowed us to use minimal versions of the PURE system that were unable to support GFP synthesis unless all genes coding for the translation proteins were included in the DNA brush. The composition of the minimal version of the PURE system that kick-started the reaction was pre-determined empirically using a deletion recovery assay for each and every one of the translation proteins and for their subgroups in bulk solution reactions, allowing us to define the transition point to cell-free biogenesis of the entire translation machinery, excluding ribosomes.

## Results

### A delta-PURE recovery assay

Of the thirty-six proteins constituting the PURE system, we focused on the minimal set of proteins that are directly involved in protein synthesis (Fig. 1b). The remaining proteins are either contributing indirectly through energy regeneration or are redundant (e.g. release factor 2, as we only use the ochre stop codon). The thirty translation proteins can be divided into 4 subgroups, defined by their biochemical functionality: twenty aaRSs, each of which covalently charges a tRNA with its corresponding amino acid; four initiation, three elongation and three termination factors, to a total of ten proteins involved in the three stages of the translation cycle (Fig. 1b).

We first verified that each of the translation proteins was synthesized from its gene in the PURE system (Supplementary Fig. S1). Then, to probe the functionality of each translation protein coupled to its synthesis within the reaction, we designed a deletion recovery assay in which a PURE system is prepared without the protein of interest (P_i_), creating a ΔPi-PURE (Fig. 1a). To measure the effect of P_i_ deletion on gene expression yield and rate, we added a reporter gene coding for GFP and observed the GFP fluorescence intensity time traces, FI(t). The addition of DNA coding for P_i_ to the ΔP_i_-PURE would indicate if and to what extent the nascent protein could compensate for its initial deletion and recover the PURE reaction.

Being a minimal system, all components in the PURE system are essential, and recovery is anticipated only if there is some trace amount of each Pi to kick-start its own synthesis. We tested the intrinsic activity level of all the different ΔPi-PURE versions via the expression of a reporter gene and found that it was highly dependent on the purity of the ribosomes in the reaction (Supp Fig. S2). Depending on the method of purification, different sources of ribosomes greatly changed the GFP synthesis activity of ΔPi-PURE versions, often reaching levels close to a full PURE system which would mask any recovery due to nascent Pi. We chose to continue with the commercial preparation of ribosomes (Methods) with the overall lowest activity of ΔPi-PUREs, to establish the minimal PURE composition required to kickstart biogenesis.

### Self-synthesis and functionality of PURE-synthesized aaRSs

We started our search for the minimal PURE reaction that could kick start self-biogenesis with the group of twenty aaRSs. We set twenty parallel ΔPi-PURE solution reactions, each testing the recovery of a different aaRS by supplementing the reaction with its respective genes. Ten of the aaRSs could be recovered with the aaRSi gene only (Arg, Asn, Cys, Glu, His, Lys, Met, Ser, Thr, Trp, Tyr), while the others required the addition of the corresponding purified aaRS as a seed (Val, Ala, Asp, Gln, Gly, Ile, Leu, Pro, Phe) (Fig. 1d, e, Supp Fig. S3-8). By analyzing the change in the GFP synthesis rate in response to DNA or seed concentrations, we could establish that the nascent aaRSs were actively participating in the PURE reaction. For example, the addition of ValRS genes to ΔValRS-PURE could not recover GFP synthesis unless purified ValRS was added. We titrated the purified ValRS into ΔValRS-PURE until the GFP yield of a full PURE system was reached, determining a dynamic range between 0.25 to 5 nM (Supp Fig. S4). Within this concentration window we observed a boost in the GFP signal upon supplementing with the ValRS gene. This suggests that purified ValRS acted as a seed that was necessary to kick-start the reaction but was not sufficient to boost full recovery (Fig. 1d, top row). The increase of GFP synthesis rate due to the addition of ValRS gene indicated that nascent ValRS was actively supporting GFP translation. A ValRS seed beyond 2.5 nM was not limiting the reaction and the recovery was no longer dependent on the addition of ValRS genes (Supp Fig. S4).

For the ten aaRSs, which did not require a seed for recovery, we found that the maximal GFP synthesis rate (max. rate) increased with the corresponding aaRS gene concentration, implying that nascent aaRSs were actively supporting protein translation (Supp Fig. S7-8). Beyond a certain gene concentration, often around 1 nM, the max. rate decreased again, which could be attributed to a ‘competition’ effect, where increasing the DNA of one gene reduces the expression of another gene (GFP), most likely due to the sharing of resources^20^.

We detected an inflection point of the GFP synthesis rate in the PURE system at about 20 to 30 minutes, even for ΔaaRS_i_-PURE before supplementing with aaRS genes, indicating intrinsic GFP synthesis before recovery (Fig. 1d, bottom row). The presence of aaRS genes led to the appearance of a second peak after a significant delay, which grew in intensity and emerged faster with increasing seed concentrations. The second peak also appeared faster with higher aaRS_i_-gene concentrations (Supp Fig. S7). We attribute the second peak to the activity of nascent aaRS_i_ contributing to its self-synthesis and GFP synthesis. We interpret the difference between the two peaks to be indicative of the time required to relieve the bottleneck created by a lack of aaRS_i_, stemming from the time it takes to synthesize sufficient aaRSi to aminoacylate its tRNAs. Thus, we identified a seed and gene concentration for all twenty aaRSs to sufficiently kick start the reaction yet still be contingent on aaRS synthesis and activity, thereby demonstrating that each was functional upon its synthesis in the PURE system (Fig. 1e, Supp Table S1). We found that when a seed was required, on average, it was between 1-2% of the concentration of the aaRS in the full PURE, often reaching sub-nanomolar concentration, with GlyRS being the exception, requiring a seed of 42%.

We defined a recovery score as the difference between the max. rates of ΔPURE reactions with and without aaRS genes, expressed as a percentage of the max. rate of a full PURE. We looked for correlations between the calculated recovery score of each aaRS and the characteristics of the reaction components, such as kinetic rate constants, size of protein or substrate amino acid, hydrophobicity of amino acids or tRNA abundance^21^. We found a negative correlation between the length of the aaRS gene and the recovery score (Kendall’s τ = −0.42, Supp Fig. S9) and between the length of the aaRS gene and the possibility to recover without the need for an external seed.

### Co-synthesis and recovery of twenty aaRSs by DNA brush localization

We attempted a recovery assay for all twenty aaRSs simultaneously by assembling a PURE solution reaction seeded with aaRSs proteins according to their recovery score and supplemented with all their genes. No GFP signal was detected above the background, implying that co-synthesis yields of twenty aaRSs were not sufficient for the self-biogenesis of an active PURE (Supp Fig. S10). To overcome this capacity barrier, we immobilized the DNA coding for all the twenty aaRSs on a glass surface as mixed DNA brushes at high density^22^ (Fig. 1f). The DNA brushes included in addition genes coding for reporter GFP fused to a Hemagglutinin tag (GFP-HA). The surface surrounding the DNA brushes was patterned with HA-specific antibodies for trapping nascent GFP on the surface for sensitive detection by TIRF imaging. We immobilized the DNA as triplet brush clusters on the same glass surface at a sufficient distance to avoid crosstalk by diffusion throughout the experiment. Each cluster had a constant 2% GFP-HA gene fraction and a different aaRSs gene fraction. To keep a constant gene expression rate, the overall DNA density was kept constant by supplementing brushes with DNA coding for an unrelated gene^23^.

Upon addition of a PURE reaction seeded according to the recovery assay of the aaRSs, a GFP fluorescent signal accumulated on the surface traps in surface areas surrounding brushes (Fig. 1f-h, Supplementary Figs. S10-12), increasing in time and in response to the fraction of aaRSs genes in the brushes. We concluded that the localization of the PURE machinery at the DNA brushes facilitated the localization of all the nascent aaRSs in the vicinity of the brush at sufficient concentrations to drive GFP synthesis^19^, thus demonstrating simultaneous recovery of twenty aaRSs in one PURE reaction.

### Self-recovery of elongation factors

Attempting to demonstrate recovery of the EFs, we recalled that their corresponding ΔPi-PURE reactions had no GFP signals (Supp Fig. S2) and that their concentration in the PURE system is in the micromolar range, in comparison to nanomolar for the aaRS. Accordingly, we had to scan a larger dynamic range of seed concentrations until full gene expression activity was recovered (Ef-Ts ∼400 nM, EF-G∼100 nM, EF-Tu∼1800 nM, Supp Fig. S12-13) compared to the aaRS recovery (e.g. ValRS,∼3 nM, Fig. S4). Next, for both EF-Ts and EF-G, the addition of the respective gene increased the GFP yield, the max. rate and the kickoff time, demonstrating the functionality of nascent EF-Ts and EF-G (Fig. 2b, c, Supp Figs S12). For EF-Tu, at all tested seed concentrations, the addition of the EF-Tu gene reduced GFP synthesis rather than boosting it (Supp Fig. S13). The fact that we could not find conditions for EF-Tu recovery is reflected in the calculated recovery score, which was close to zero for EF-Tu, 25% for EF-G and 50% for EF-Ts, (Fig. 2d).

**Figure 2.**
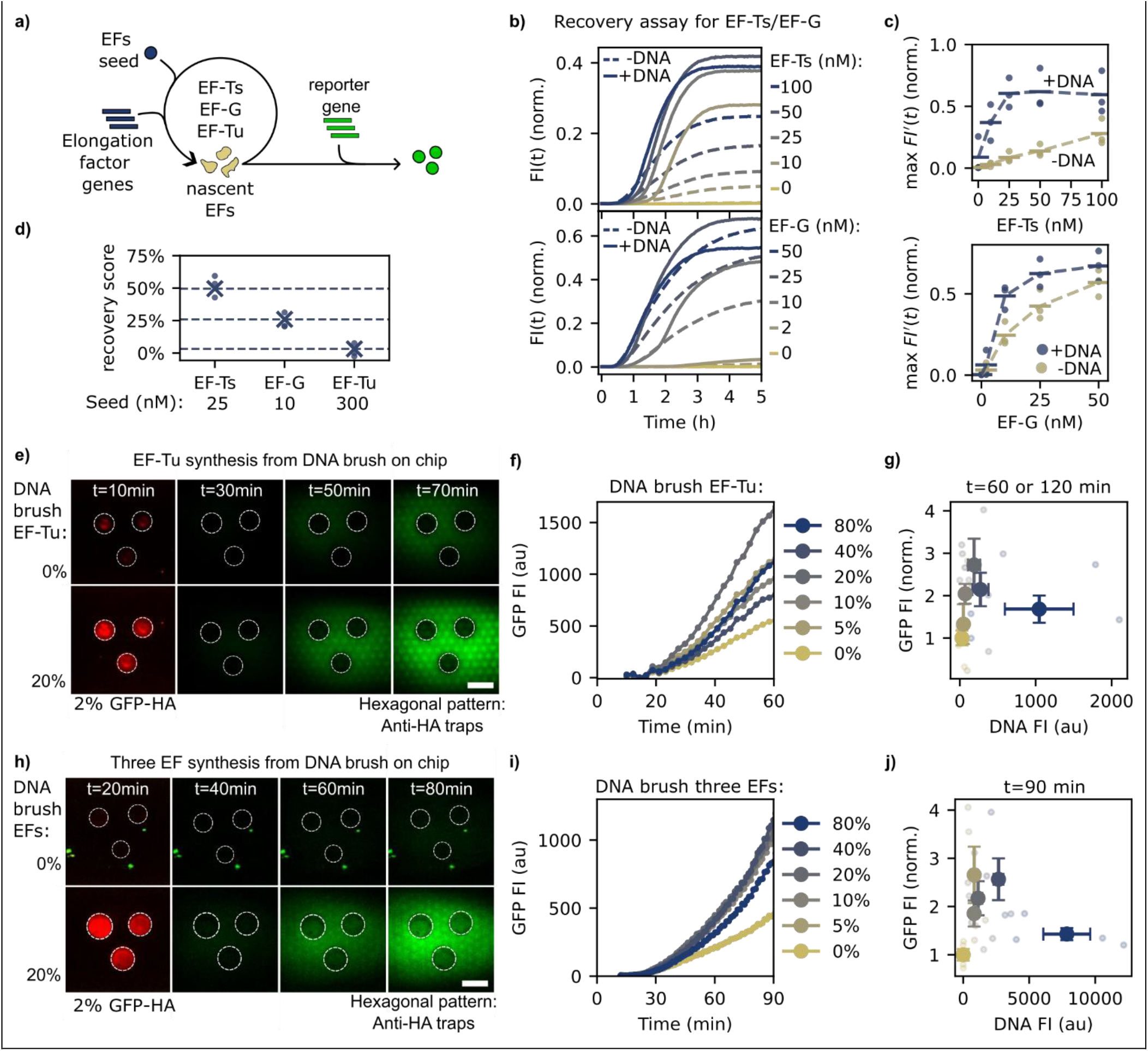
Biogenesis of three EFs from DNA brushes. **a)** Scheme: self-synthesis of the three EFs from their genes in a seeded ΔEF-PURE system with a reporter gene. **b-d)** Recovery assay in a microplate well. **b)** FI(t) of ΔEF-Ts-PURE (top, solid lines with [DNA_EF-Ts_] = 1 nM.) with different amounts of purified EF-Ts and of ΔEF-G-PURE (bottom, solid lines with [DNA_EF-G_] = 0.75) with different amounts of EF-G. [DNA_GFP_] = 1 nM. FI(t) are normalized to the max. int. of a full PURE sample. **c)** Max. rate of GFP FI(t) from ΔEF-Ts-PURE as a function of the EF-Ts seed (top) and ΔEF-G-PURE as a function of the EF-G seed (bottom). Max. rate is normalized to the max. rate of a full PURE sample. Mean values (horizontal markers) for n=3. **d)** Recovery scores derived from recovery assays with seed concentrations that yielded the highest score for each of the three EFs. **e-g)** TIRF images of self-synthesis of EF-Tu from surface-immobilized DNA brushes, marked as in Fig. 1f. **e)** Top row: DNA brushes code for a non-interacting control protein (DNA not labeled), bottom row: DNA brushes code for EF-Tu (DNA-labeled). All brushes contain 2% GFP-HA genes. Upon exposure to a PURE with a seed of [EF-Tu]=1000nM, surface-bound GFP-HA accumulates around DNA brushes dependent on the fraction of EF-Tu genes. **f)** average GFP FI(t) from TIRF microscopy images for DNA brushes with different fractions of EF-Tu genes. **g)** Normalized mean GFP FI and its s.e.m. as a function of the DNA FI (EF-Tu + GFP DNA) for different DNA brushes, n=4. The selected time points for the GFP level vary between t=60 or t=120 min, depending on the expression kinetics (Fig. S14). **h-j)** Synthesis of all three EFs from DNA brushes. **h)** Images are labeled as in subfigure e), but with all three EF genes in the DNA brush, with a gene ratio [Ts]:[G]:[Tu] of 1:2:3 ratio. The seeded ΔEF-PURE system [EF-Ts]=100nM, [EF-G]=25nM, [EF-Tu]=1000nM. **i)** Average GFP FI(t) from TIRF microscopy images for DNA brushes with different fractions of EF genes. **j)** Normalized mean GFP FI at t=90 min and its s.e.m. as a function of the DNA FI (EFs + GFP DNA) for different DNA brushes. n=4. For g) and j) values are normalized to the average GFP Fl of a DNA brush with 0% EFs genes. All scale bars = 200 μm.

The lack of recovery via EF-Tu expression could be caused by its extremely high concentration requirement in the PURE system for efficient protein synthesis^24^. To rule out the possibility that the lack of recovery was not due to a nonfunctional nascent EF-Tu, we separated the recovery reaction into two steps (Methods). In the first step, we added EF-Tu-DNA to a seeded PURE reaction, and after incubation, allowing for sufficient EF-Tu synthesis to occur, we transferred a small fraction of this reaction to a fresh ΔEF-Tu-PURE reaction, including the reporter gene (Methods, Supplementary Fig. S13). We observed that the GFP signal in reaction 2 increased with the EF-Tu gene concentration in reaction 1, thus confirming that nascent EF-Tu was functional and that the recovery was most likely not efficient enough to detect the activity of nascent EF-Tu in one step.

To demonstrate the biogenesis of EF-Tu in a single reaction, we immobilized EF-Tu DNA on a glass surface in a triplet DNA brush cluster, similar to the aaRSs experiment (Fig. 2e, 1f, respectively), with every cluster having a different fraction of EF-Tu DNA, from 0 to 80%, and a 2% constant fraction of GFP-HA genes. Upon addition of a ΔEF-Tu-PURE reaction seeded with 1 micromolar purified EF-Tu, we observed over the course of one-hour accumulation of GFP-HA trapped on the surface in the vicinity of DNA brushes, demonstrating that the EF-Tu synthesized from the brushes was functional in the PURE reaction (Fig. 2e, Supp Fig. 14). The GFP-HA signal increased in response to larger fractions of EF-Tu-DNA up to 40%, with a slight decrease at 80% most likely due to resource competition.

Encouraged by the self-recovery of EF-Tu when encoded in DNA brushes, we attempted simultaneous self-recovery of all three elongation factors coded in the same DNA brushes. Upon addition of a ΔEFs-PURE seeded with EF-Ts, EF-G, and EF-Tu at concentrations determined by the preliminary solution and surface experiments, GFP-HA signals accumulated on the surface surrounding DNA brushes in response to the increase of EFs genes fraction, with a maximum GFP signal at 40% and a sharp drop at 80% (Fig. 2f, Supp fig S14).

### A group recovery assay for nascent Initiation and Release factors

We attempted to demonstrate self-recovery of Initiation and Release factors (Fig. 3a, d) even though single deletions of each of the initiation group (IF1, IF2, IF3, MTF) and of the release group (RF1, RF3 and RRF) did not create a significant bottleneck for GFP synthesis due to intrinsic seeds (Supp Fig. S2, S15). To create a bottleneck that restricts GFP synthesis, we assembled ΔP_i_-PURE solutions with all deletion permutations of the four- and three-member initiation and release groups, respectively. All combinations, except for Δ(IF1+MTF), had a lower max. GFP synthesis rate than a full PURE reaction that increased upon the addition of the corresponding genes, indicating that the nascent proteins were synthesized and active in protein translation (Fig. 3b,e, Supp Fig. S15). The Δ(IF1+MTF) exception is consistent with the fact that these factors were shown to be the least essential for protein synthesis *in vitro* ^3^. Nevertheless, by calculating the recovery score of each of the combinations, we observed an increase in the recovery score of Δ(IF1+IF3) and Δ(IF3+MTF) compared to their single deletion scores (Fig. 3c,f), indicating functionality of all nascent components. This conclusion is supported by the observation that the recovery score of the tri- and tetra-deletions Δ(IF1+2+3) and Δ(IF1+2+3+MTF) respectively, had an increase in the kickoff time in the presence of the DNA (Fig. 3c). That is, relieving the bottleneck created by the lack of four proteins took about eight times longer (40 min) compared to the lack of one (5 min). We concluded from these recovery assays in bulk solutions that all seven nascent Initiation and Release factors were participating in GFP synthesis upon their self-synthesis in the PURE reaction.

**Figure 3.**
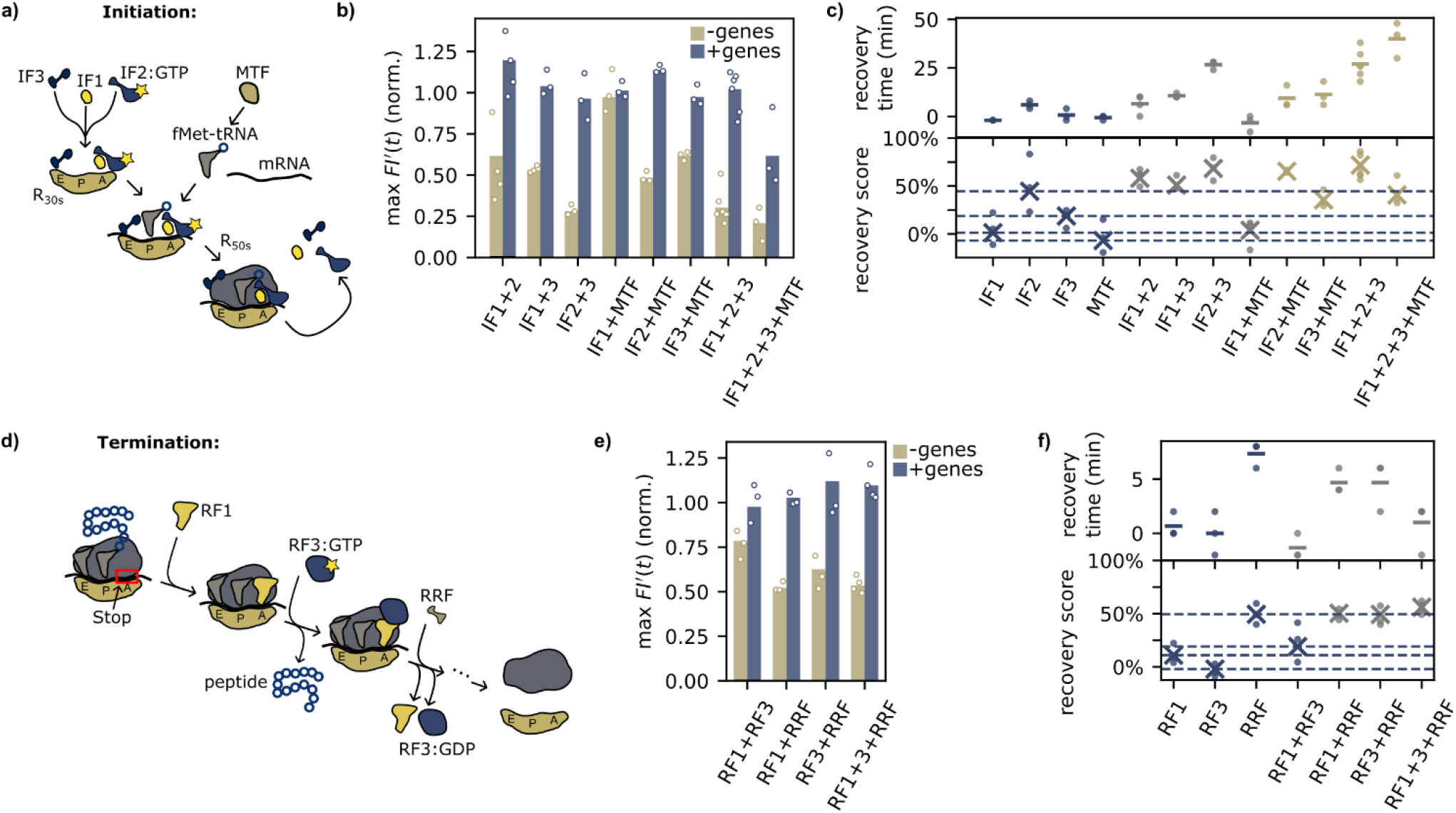
Group recovery assay for IFs and RFs. **a)** Scheme: translation initiation process involving the three initiation factors IF1, IF2 and IF3 and the methyltransferase (MTF). **b)** Bar plot of the normalized max. rate of ΔPURE for all the permutations of the three IFs and MTF +/−their corresponding genes. For each gene: [IF-DNA] = 0.1 nM and [GFP-DNA] = 0.4 nM. PURE recovery measurements in a microplate well with min. n=3. **c)** Bottom: Recovery scores derived from the recovery assays of each IFs and the subgroups from b). Dotted horizontal lines at the value of the individual protein scores as a guide to the eye for comparing the subgroup scores. Top: corresponding recovery times. **d)** Scheme of the translation termination process involving the two release factors RF1 and RF3, as well as the ribosome recycling factor RRF. **e)** Bar plot of the normalized max. rate of ΔPURE for all the permutations of a grouping of RF1, RF3, and RRF +/−their corresponding DNA. For each gene: [RF-DNA] = 0.1 nM and [GFP DNA] = 0.4 nM, minimum n=3. **f)** Bottom: recovery scores for the recovery assay from e). Top: corresponding recovery times.

### Recovery of a stringent PURE reaction by self-synthesis of all auxiliary translation proteins

Having support for the self-recovery of each of the auxiliary protein components and their subgroups, we attempted the co-recovery of all of them simultaneously in a single PURE reaction (Fig. 4a). We immobilized genes coding for all IFs, EFs, RFs, and aaRSs as a mixed DNA brush on a glass surface (Fig. 4b, c), with relative gene fractions among the PURE auxiliary proteins’ genes based on the recovery scores from the preliminary experiments. Clusters of DNA brushes on the same surface had variable fractions of the PURE auxiliary proteins’ genes, ranging from zero to 90%, keeping constant the GFP-HA fraction at 2% and the overall DNA density by supplementing brushes with DNA coding for an unrelated gene.

**Figure 4.**
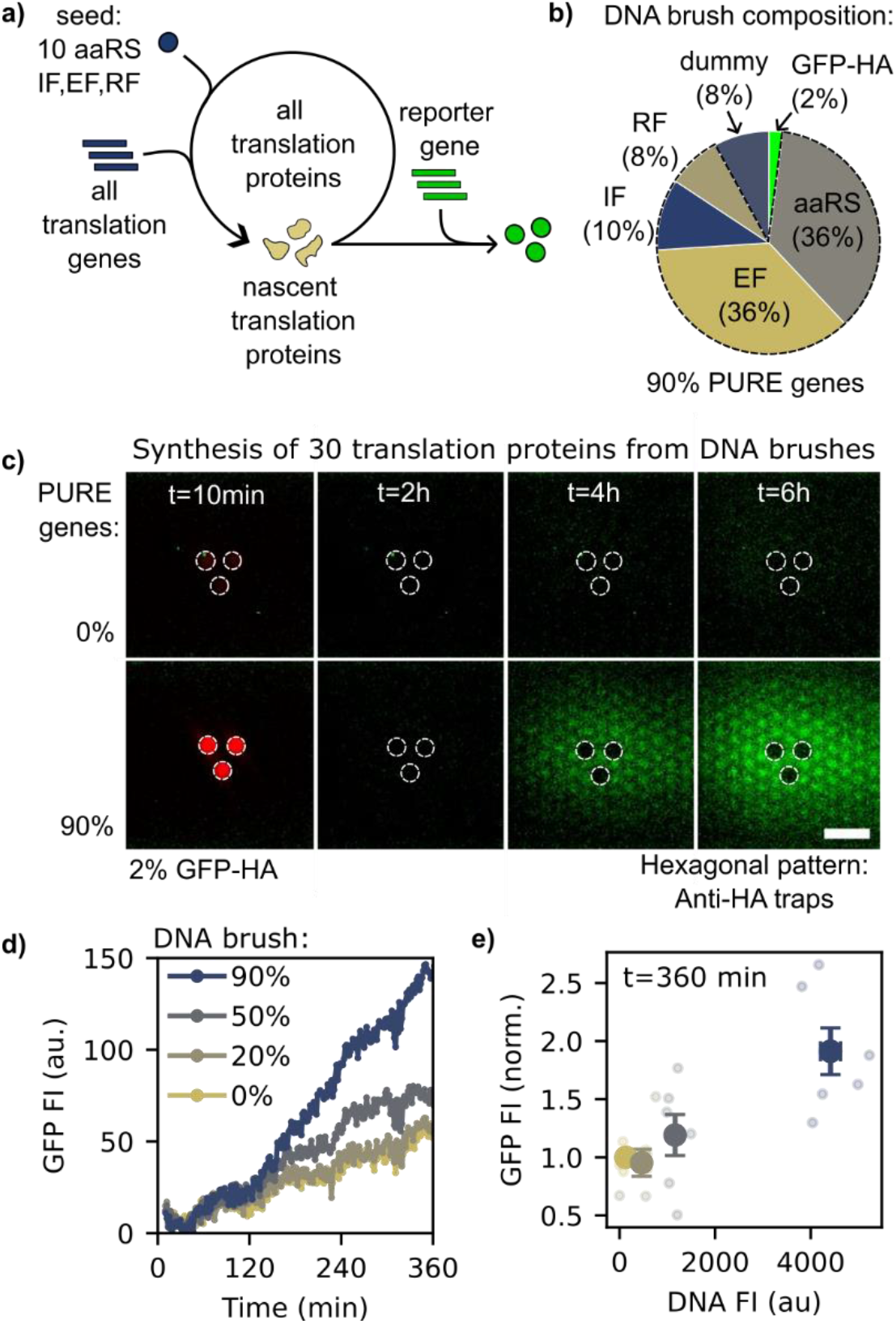
Biogenesis of all translation proteins from DNA brushes. **a)** Scheme: recovery assay for all translation factors from a minimally seeded PURE system (see Table S2). **b)** Composition of the DNA brush. **c)** TIRF microscopy images of synthesis of all translation proteins from surface-immobilized DNA brushes, marked as in Fig. 1f. Top row: DNA brushes code for a non-interacting control protein (DNA not labeled). Bottom row: DNA brushes code for all translation genes (DNA-labeled). All brushes contain 2% GFP-HA genes. Upon exposure to a seeded ΔPURE, surface-bound GFP-HA accumulates around DNA brushes depending on the fraction of translation genes. Scale bar = 200 μm. **d)** Average GFP FI(t) for DNA brushes with different fractions of translation genes. **e)** Normalized mean GFP FI at t=6 h and its s.e.m. from TIRF microscopy as a function of the DNA FI (translation protein-DNA + GFP DNA) for different DNA brushes. n=6. Values are normalized to the average GFP Fl of a DNA brush with 0% translation genes.

The addition of the most stringent PURE reaction, seeded only for aaRS and EFs, did not increase GFP expression in the vicinity of all genes (Supp Fig. S16). Only with the addition of a seed for the IF and RFs did GFP-HA accumulate on the surface in response to an increase in the PURE genes (Fig. 4c,d,e, Supp S17, Table S2). The long recovery time was consistent with the many barriers that had to be overcome in the PURE reaction involving parallel synthesis of thirty different translation proteins, each required for the synthesis of the other. Thus, the localization of the DNA brushes was essential to increase the apparent concentration of all nascent proteins, attracting them to their source genes and participating in the synthesis of GFP.

## Conclusion

We showed that the thirty proteins constituting the translation machinery of the PURE system, excluding the ribosome, could be synthesized simultaneously from their corresponding genes by the PURE reaction itself, thus laying the foundation for a self-regenerating PURE system. While some of the translation proteins or even subgroups could self-recover in a PURE reaction in solution, the entire translation machinery was dependent on high-density immobilized DNA to create localization conditions essential for multi-self-recovery.

We demonstrated in detail how deletions of each translation factor affect the gene expression rates in our PURE system. Using a recovery assay, we studied the recovery process of gene expression for each protein and found the limitations in terms of the PURE gene concentration. We could follow the recovery process’s kinetics and identify the time it takes for the newly synthesized protein to release the bottleneck imposed by the ΔPURE, which varies considerably from protein to protein. By screening different seeding concentrations in the ΔPURE, we determined the minimal concentration for each protein to have a full-functioning PURE, and how sensitive it is to its fluctuations of concentrations in that regime. We observed that seeding is essential for the recovery of gene expression and how, in the case of the aaRS, a high molecular weight of the proteins seems to be an obstacle to recovery without a seed. These results open avenues towards a self-sustaining PURE system that can self-regenerate its components, replacing the purified components with synthetic ones.

## Materials and Methods

### PURE system preparation

#### Protein purification

The PURE system was prepared according to Shimizu *et al*. ^25^. *E. coli* BL21(DE3) *pLyS* strain was used for protein expression. All plasmids encoding PURE proteins were obtained from Y. Shimizu. Plasmids were transformed and amplified in *E. coli* DH5α, except for plasmids coding for AlaRS, AsnRS, ThrRS, IF1-3, EF-G, RRF, CK, T7RNAP, PheRS, which were transformed to *E. coli* JM109 due to their expression vector (pQE30). All cultures were grown at 37 °C, 250 rpm. Overnight cultures were grown in 5 mL of LB medium (LBX0102, Formedium, UK) with 100 μg/mL of ampicillin or 50 μg/mL of kanamycin. Each strain was inoculated in a flask with 1 L of LB and grown to OD_600_ = 0.6before induction with 0.1mM of isopropyl β-D-1-thiogalactopyranoside (IPTG from Applichem, Germany) for 3 h, and then harvested after centrifugation and stored at −80 °C.

Protein purification buffer compositions are as follows. Buffer A: 50 mM HEPES-KOH (8042391, Bio-Lab, IL) pH 7.6, 1 M ammonium chloride (3384-12, Avantor, USA), 10 mM MgCl_2_ and 7 mM 2-mercaptoethanol (b-ME, AlfaAesar, ThermoFisher Scientific, USA, only added immediately before use). Buffer B: 50 mM HEPES-KOH pH 7.6, 100 mM KCl (7447-40-7, Merck, DE), 10 mM MgCl_2_, 500 mM imidazole (GB9580, Glentham Life Sciences, UK) and 7 mM b-ME. HT Buffer: 50 mM HEPES-KOH pH 7.6, 100 mM KCl, 10 mM MgCl_2_, and 7 mM b-ME. Protein storage buffer: 50 mM HEPES-KOH pH 7.6, 100 mM KCl, 10 mM MgCl_2_, 30% glycerol (007120223300, Bio-Lab, IL) and 7 mM b-ME.

The cell pellets were re-suspended in 30 mL of buffer A and lysed by sonication for 3 min on ice (Sonics Vibra-Cell VC 750; probe tip diameter: 13 mm; 20 s:20 s pulse; 70% amplitude). Cell debris was removed by centrifugation (25,000 cf, 30 min, 4 °C). The protein was then purified from the supernatant either on a gravity-flow column or with an FPLC (Äkta avant, Cytiva, USA). For the gravity-flow method, the supernatant was mixed with 2–3mL of equilibrated resin and incubated for up to 2 h at 4 °C. After the incubation, lysate was loaded on a column (Econo-Pac, Bio-Rad, USA). The column was washed with 30 mL of a wash buffer (95% buffer A, 5% buffer B) and eluted with 15 mL of an elution buffer (10% buffer A, 90% buffer B). For the FPLC method, the supernatant was loaded onto a 5 mL HisTrap FF Crude column (Cytiva, USA), the column was then washed with 20 column volume of wash buffer (95% buffer A, 5% buffer B). Proteins were eluted by a linear 20 column volume gradient (5 to 100% buffer B) with a flow rate of 1 ml/min. Following either method, the elution fraction was dialyzed twice against HT buffer followed by once against protein storage buffer. Protein purity was assessed with SDS-PAGE gel (4-20% GeBaGel, Gene Bio-Application, IL) and Coomassie staining (InstantBlue, Abcam, UK). The protein solution was concentrated using a Vivaspin 20 centrifugal concentrator (molecular cut-off depending on the protein, Cytiva, USA). Protein concentrations were estimated by absorbance at 280 nm using calculated protein extinction coefficients. Proteins were then stored at −80 °C in aliquots.

#### Ribosome purification

Ribosomes were purified from the *E. coli* strain *A19* (no antibiotics) after lysis using a French press (Constant Systems Cell Disruptor, England) following three different published protocols (Fig. S2): Anion Exchange Chromatography, as described in Trauner *et al*.^26^ using a monolithic column (CIMmultus QA 80ml, BIA separations) connected to an FPLC (Äkta avant, Cytiva, USA); Hydrophobic Interaction Chromatography followed by ultracentrifugation was performed according to Shimizu *et al*^24,25^ with a 5 mL HiTrap Butyl FF column (Cytiva, USA) connected to an FPLC (Äkta avant, Cytiva, USA). The ribosomes were layered on a sucrose cushion buffer (10 Tris, pH =8, Mg(OAc)_2_ 14mM, KOAc 60mM, Sucrose 1.1M, b-ME 6mM) and pelleted using a using Ti-70 rotor at 55,000 rpm, 4°C for 17 hours. For the third purification method, we followed the protocol of Halfon *et al*.^27^, we first used an ultracentrifuge with a sucrose cushion buffer like previously mentioned, followed by sucrose gradient ultracentrifugation on a 10%-40% sucrose gradient with the same buffer 10 Tris, pH =8, Mg(OAc)_2_ 14mM, KOAc 60mM, b-ME 6mM) for 16 hours at 19000 RPM in a swing SW-28 rotor. Ribosome fractions were collected according to A260 values and pelleted at 45,000 rpm using a Ti-70 rotor for 19h. After purification, the ribosomes from each method were dissolved in or dialyzed against 1x Ribosome buffer (20 mM Hepes, pH=7.6, 30 mM KCl, 6 mM Mg(OAc)_2_, 7 mM 2-mercaptoethanol) and stored in aliquots at −80 °C.

#### Buffer preparation

PURE buffer was prepared by stepwise addition of the following solutions, in the order they are listed here, starting with the volume of water necessary to reach the desired concentrations. The final concentrations are listed in the PURE assembly below. The concentrations listed here are for a 2X buffer stock: 100 mM HEPES-KOH pH 7.6 200 mM K-glutamate, 20 mM Mg(OAc)_2,_ 4 mM ATP, 4 mM GTP, 2 mM CTP, 2 mM UTP, 108 units of A280/mL tRNA, 40 mM creatine phosphate, 2 mM DTT, 20 μg/mL folinic acid, 4 mM spermidine, 0.6 mM each amino acid (prepared according to ref ^28^). The solution was aliquoted and stored at −80°C. The optimal Mg(OAc)_2_ for GFP expression was determined by doing an initial Mg(OAc)_2_ scan between 10-30 mM and varied slightly between 16-22 mM between buffer preparation batches.

#### PURE assembly

The PURE system was mixed on ice. The protein solution contained (following Shimizu *et al*.^25^): 704 nM AlaRS, 30 nM ArgRS, 408 nM AsnRS, 119 nM AspRS, 23 nM CysRS, 59 nM GlnRS, 230 nM GluRS, 85 nM GlyRS, 17 nM HisRS, 381 nM IleRS, 41 nM LeuRS, 109 nM LysRS, 27 nM MetRS, 134 nM PheRS, 154 nM ProRS, 38 nM SerRS, 83 nM ThrRS, 29 nM TrpRS, 12 nM TyrRS, 16 nM ValRS, 1.036 μM IF1, 405 nM IF2, 456 nM IF3, 636 nM EF-G, 2.255 μM EF-Tu, 1.588 μM EF-Ts, 247 nM RF1, 162 nM RF3, 458 nM RRF, 568 nM MTF, 89 nM CK, 64 nM NDK, 100 nM T7 RNA P., 132 nM MK, 30 nM PPiase; The buffer and energy solution (see Buffer Preparation). The ribosomes: 2 μM of commercial (Solution III of PUREfrex 2.1, Gene Frontier, JP) or purified ribosomes. We added to each reaction 1.25%(v/v) GroE mix Gene Frontier, JP) and 2.5%(v/v) DnaK mix Gene Frontier, JP). PURE assembly was mostly done by hand pipetting. Seed concentrations of single PURE proteins were diluted by hand in protein storage buffer and then dispensed with a liquid handling robot (iDOT, Dispendix Germany) into a premixed ΔP_i_-PURE. For the ribosome purity test (Fig. S2), we pre-mixed solutions containing 5 proteins by hand, and then we assembled the PURE with an Echo 525 (Labcyte, USA).

### DNA preparation

For expression in PURE, all translation genes were placed under a T7 promoter and a strong ribosome binding site (RBS). All genes were cloned into pET-plasmids using Gibson Assembly (NEBuilder HiFi Assembly Master Mix, E2621, NEB, USA). Plasmids were amplified in *E. coli* DH5α and purified using Wizard SV-Gel Miniprep (Promega, USA), and DNA concentrations were determined using a NanoDrop (NanoPhotometer, Implen, USA). Linear double-stranded DNA fragments were amplified from plasmids with Polymerase Chain Reaction (PCR) with KAPA HotStart ready mix (07958935001, Roche, Switzerland), using a reverse primer conjugated to biotin as previously described^29^ and a forward primer that was either unmodified or conjugated to an ATTO 647 fluorophore on its 5’ end (Integrated DNA Technologies, USA), and purified with the Wizard SV-Gel and PCR Clean-Up System (Promega, USA).

For surface immobilization, DNA was mixed with streptavidin (S4762, Sigma-Aldrich) at a 1.4:1 streptavidin: DNA ratio in 1x phosphate-buffered saline (PBS, 02-023-5A, Sartorius, Germany) and 7% glycerol (Bio-Lab, Israel), forming a DNA-streptavidin conjugate. The correct amplification of DNA fragments and DNA-streptavidin conjugates was verified with 1% agarose gel electrophoresis. The solution used to create DNA brushes contained various concentrations of the genes of interest, as indicated in Table S2. The final DNA: streptavidin concentration for a DNA brush solution was 150 nM.

### Bulk solution experiments

Bulk expression experiments in the PURE system were conducted at 37 °C in a ClarioStar plate reader (BMG Labtech, Germany). DNA was added to the PURE reaction prepared as described above. Volumes of 10 μL were pipetted in a black optical-bottom 384-well plates (Nalge Nunc International, USA), sealed with a sticky transparent foil (SealPlate, Excel Scientific, USA), and spun down at 1,000 rcf for 30 seconds. GFP fluorescence was measured with an excitation filter of 470/15 nm, a dichroic filter of 491 nm, and an emission filter of 515-20 nm. Every experiment contained a well with full PURE with the same GFP-DNA concentration for normalization purposes between experiments. For repeats of the same experiment, the critical components of the PURE reaction, such as the protein seed or the gene DNA, were always added separately to each reaction.

### Gel electrophoresis analysis of fluorescently labeled nascent proteins

A more concentrated PURE protein and energy/buffer solution was mixed following the composition provided by PURE version 2 by Kazuta *et al*. ^10^ Commercial ribosomes (Solution III of PUREfrex 2.1, Gene Frontier, JP) were used according to their published manual. For each PURE gene, 1 nM of DNA and 4%(v/v) Green-Lys tRNA (FluoroTect Green_Lys_, Promega, USA) were added to the assembled PURE and incubated for 4h at 37 °C in a PCR thermocycler (Bio-Rad, USA). The reactions were incubated for 2 minutes at 60 °C and assessed with SDS-PAGE (160V 1h) gel (4-20% GeBaGel, Gene Bio-Application, IL) and imaged with a Typhoon FLA 9000 (GE, USA).

### Glass surface experiments

#### Slide preparation

Fused-silica slides (24×24×1mm, UQG Optics, UK) were coated with a biocompatible photosensitive monolayer^30^ of a polymer formed by a polyethylene glycol backbone with a protected amine and a triethoxysilyl group at either end. The slides were first cleaned in boiling ethanol (830109326, 96%, Gadot Group, IL) for 10 minutes, followed by base piranha cleaning (1:1:4 H_2_O_2_ (000855032300, Bio-Lab, IL):NH_3_ (105432, Merck, DE):H_2_O) at 70°C for 10 minutes. The slides were incubated for 20 min with a 1 mg.ml^-1^ concentration of polymer dissolved in dried toluene (244511, Sigma-Aldrich, USA), then rinsed with toluene (Bio-Lab, Israel) and dried. Surface amines were then deprotected with UV exposure (UV-KUB, FR, 365 nm, 2.5 J/cm^2^) through a custom photomask with an array of 30 μm hexagons (CAD/Art Services. USA). The slides were immediately incubated with 0.5 mg/ml biotin 3-sulfo-*N*-hydroxysuccinimide ester (EZ-link NHS biotin, 20217, Thermo Fisher Scientific, USA) in 0.2 M borate-buffered solution pH 8.6 (Thermo Fisher Scientific) for 30 min, then rinsed with water and dried. The slides were then fixed on custom fused silica prisms (Zell Quarzglas und Technische Keramik, Germany) with Frame-Seal Slide Chambers adhesive (Bio-Rad, USA). Frame-Seal Slide Chambers of size 15×15 mm (vo=65ul) were applied to the polymer-treated side of the fused-silica slides to act as reaction chambers for the DNA brushes.

#### DNA deposition

The biotinylated surfaces of the wells were patterned with nano-liter DNA-SA droplets using the GIX Microplotter II (Sonoplot Inc., USA), and incubated overnight at room temperature and 50-65% humidity to allow DNA brush build-up through biotin-streptavidin binding. We defined 12 positions for DNA brushes on each chip, ensuring at least a 3 mm difference between positions. We spotted three different identical DNA brushes at each position to increase reproducibility (also referred to as a cluster) and increase GFP signal strength. In each experiment, we titrated the concentration of the PURE genes in the DNA brush cluster in question (e.g. 0%, 20%, 50%, 90%). The dilutions of the PURE genes were done with a control gene that codes for a non-interacting protein so that the total DNA concentration remained at 150 nM. On each glass slide, we repeated the titration series of DNA brush clusters at least twice.

#### Capture antibodies immobilization

Biotinylated anti-HA antibodies (50 μg/ml, High Affinity, 12158167001, Roche, Sigma-Aldrich) were mixed with streptavidin (S4762, Sigma-Aldrich) at a concentration ratio of 1.5:1 in 1x PBS and incubated for 30min at 4 °C, after which the mix was diluted to 5 μg/ml of antibodies in 1x PBS. 200 μl of this solution was applied to the surface of the chip after the overnight DNA incubation and incubated for one hour at 4 °C. Afterwards, the surface was washed several times with 1x PBS, then with 50 mM HEPES pH7.4.

#### Protein expression from DNA brushes

The HEPES buffer was washed and exchanged with a PURE mix in four consecutive washes of 50 μL each. The well was then sealed with a glass coverslip. The slide and attached prism were immediately positioned on the microscope stage and kept at 17 °C with a temperature-controlled holder. Temperature was increased to 37 °C to start gene expression. Different experiments on a chip were repeated at least two times.

### TIRF imaging

Prior to imaging, the space between the prism and the slide was filled with index-matching liquid (Cargille, USA). The wells were imaged with a custom-made Total Internal Reflection Fluorescence (TIRF) system. An upright microscope (Olympus BX51WI, JP) was equipped with a motorized stage (Scientifica, UK) and a laser system for TIRF excitation. Two lasers (OBIS 488-150 LS and OBIS 647 LX, Coherent, USA) were coupled into a single-mode optical fiber (Oz optics, CAN), after which the beam was collimated and directed to the prism with a goniometer (Thorlabs, USA) at an angle of total internal reflection on the surface of the chamber, creating an evanescent wave inside the chamber. Images were acquired using an Andor iXon Ultra camera (Andor Technology, UK) and a 10x Olympus objective. The set-up was controlled through a custom-made LabView code (National Instruments, USA). The DNA signal was measured with the red laser at the beginning of the acquisition at 17 °C. The GFP signal time series was measured in the green laser afterwards at 37 °C.

### Data processing and analysis

#### Plate reader data

Automated data processing and analysis were implemented with Python v3.7, and data was generally handled with the *pandas (v*2.2.3) package. For bulk solution experiments, fluorescence intensity time traces were normalized:

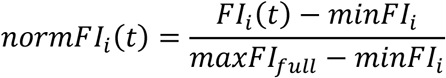

where FI_i_(t) is the fluorescence intensity time trace for sample i, minFI_i_ is the minimum fluorescence intensity value of sample i (background level), and maxFI_full_ is the maximal value (only after the signal has saturated with time ∼4-8h) of a full PURE reaction from the same experiment with the same concentration of reporter gene.

The FI(t) was smoothed by applying the Savitzky-Golay filter (imported from *scipy* (v.1.14.1) package with a window length = 7 and a polynomial order = 0) before the first rate is calculated. The normalized max. rate is calculated as the max. rate of the sample divided by the max. rate of a full PURE sample from the same experiment. The second rate is calculated by applying the Savitzky-Golay filter to the first-rate and then calculating its rate again. The normalized second rate is calculated by dividing the max. of the 2^nd^ rate of sample i, by the max. of the 2^nd^ rate of a full sample.

The recovery score is defined as the difference between the normalized max. rate in the presence of the gene and the normalized max. rate in the absence of the gene.

Kendall’s rank correlation coefficient was calculated with the *kendalltau* function with variant b in the *scipy* (v.1.14.1) package. For categorical variables (e.g. Seed: ‘Yes’ or ‘No’), the rank-biserial correlation *r* was calculated with:

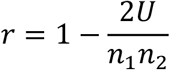

where U is calculated by the Mann-Whitney U test from the *scipy* (v.1.14.1) package, and n_1_ and n_2_ are the sample sizes of the groups that do and do not require an external seed, respectively.

#### TIRF microscopy data

For TIRF microscopy images, we averaged the GFP signal of the whole image for each time frame. The background signal was determined to be the green channel signal from one of the first 5 frames and then subtracted from the GFP signal of all other time frames. For experiments with a very low GFP signal compared to the background, we defined two regions of interest (ROI) with the same area. The first ROI-1 is nine circles positioned in the center of the hexagons that contain the antibody traps for GFP-HA (specific GFP signal), and the second ROI-2 is nine circles positioned in areas between hexagons, where the density of the antibodies is significantly reduced (background). The circle positions are chosen to be close to each other within the image to avoid differences in illumination due to the alignment of the laser. The background-subtracted GFP signal for each time frame is then calculated by the mean intensity of ROI-1 minus ROI-2.

For time series of GFP expression from DNA brushes, t=0 is set to the time point when the temperature on the prism holder reached 37 °C, determined using a temperature sensor. Image acquisition started after the focus plane had been adjusted at 37 °C. This time difference was measured and considered in the data analysis. The TIRF signal from the labeled DNA in the DNA brushes was measured by defining two ROIs of the same area, the first ROI-1, including the DNA brushes (DNA specific signal), and the second ROI-2, excluding the DNA brushes (background). The normalization of the GFP signal across glass slides for the TIRF microscopy data was conducted in the following way. For each slide, the average GFP signal from all the DNA brushes without PURE genes is calculated, and this is used to normalize the GFP signal from all other DNA brushes from the same glass slide. Then, the normalized GFP signals of DNA brushes with the same genetic content from across different slides are averaged, and the standard error of the mean is calculated.

## Acknowledgments

The authors thank André Rivalta and Ada Yonath for help with the ribosome purification, Galit Cohen and Haim Barr from the G-INCPM for help with the ECHO operation, and Ohad Vonshak for useful discussions. This research was supported by the United States Office of Naval Research (R.B.Z. and S.S.D. Award N62909-22-1-2042), and the Isak Ferdinand and Dwosia Artmann Research Fund for Biological Physics (R.B.Z.). The work of M.S.S. was supported by the Deutsche Forschungsgemeinschaft (DFG, German Research Foundation, Projektnummer 456967983). A.D. acknowledges funding from the EMBO postdoctoral fellowship, award number: ALTF 131-2020.

