## Supplementary Information for "Autonomous biogenesis of the entire protein translation machinery excluding ribosomes"

| Protein: | Protein DNA (nM): | GFP DNA (nM): | Seed (nM) | Recovery score (%) |
|---|---|---|---|---|
| AlaRS | 0.1 | 0.4 | 4.0 | 41±12 |
| ArgRS | 0.1 | 1.0 | 0.0 | 82±13 |
| AsnRS | 1.0 | 1.0 | 0.0 | 93±10 |
| AspRS | 0.1 | 0.4 | 1.5 | 73±26 |
| CysRS | 0.1 | 1.0 | 0.0 | 103±23 |
| GlnRS | 1.0 | 1.0 | 0.5 | 59±23 |
| GluRS | 0.1 | 1.0 | 0.0 | 100±40 |
| GlyRS | 0.1 | 0.4 | 35.0 | 57±37 |
| HisRS | 1.0 | 1.0 | 0.0 | 95±19 |
| IleRS | 1.5 | 1.0 | 0.5 | 16±31 |
| LeuRS | 1.0 | 1.0 | 1.0 | 37±8 |
| LysRS | 0.1 | 1.0 | 0.0 | 68±13 |
| MetRS | 5.0 | 1.0 | 0.0 | 22±2 |
| PheRS | 0.1 | 0.4 | 1.0 | 38±7 |
| ProRS | 1.0 | 1.0 | 2.0 | 43±17 |
| SerRS | 1.0 | 1.0 | 1.0 | 123±4 |
| ThrRS | 0.1 | 0.4 | 0.0 | 104±30 |
| TrpRS | 0.1 | 0.4 | 0.0 | 43±14 |
| TyrRS | 0.1 | 0.4 | 0.0 | 106±41 |
| ValRS | 1.5 | 1.0 | 1.0 | 67±14 |

**Table S1. Gene and seed concentrations for aaRS recovery assays.** Gene concentration, GFP DNA concentration and external seed concentration for the ΔaaRS$_i$-PURE recovery assays in bulk solution experiments shown in Fig. 1d. From the recovery assay, we calculated the corresponding mean recovery score and its standard deviation, for min. n=3.

| Gene | DNA conc. (nM) | Protein seed for DNA brush (nM) | Gene | DNA conc. (nM) | Protein seed for DNA brush (nM) |
|---|---|---|---|---|---|
| **AlaRS** | 6.7 | 5 | **EF-G** | 22.2 | 25.0 |
| **ArgRS** | 6.7 | 0 | **EF-Tu** | 33.3 | 1000.0 |
| **AsnRS** | 6.7 | 0 | **EF-Ts** | 11.1 | 100.0 |
| **AspRS** | 6.7 | 5 | **IF1** | 2.4 | 25.0 |
| **CysRS** | 6.7 | 0 | **IF2** | 2.4 | 25.0 |
| **GlnRS** | 6.7 | 5 | **IF3** | 2.4 | 25.0 |
| **GluRS** | 6.7 | 0 | **MTF** | 2.4 | 25.0 |
| **GlyRS** | 6.7 | 20 | **RF1** | 2.4 | 25.0 |
| **HisRS** | 6.7 | 0 | **RF3** | 2.4 | 25.0 |
| **IleRS** | 6.7 | 5 | **RRF** | 2.4 | 25.0 |
| **LeuRS** | 6.7 | 5 | | | |
| **LysRS** | 6.7 | 0 | | | |
| **MetRS** | 6.7 | 0 | | | |
| **PheRS** | 6.7 | 5 | | | |
| **ProRS** | 6.7 | 0 | | | |
| **SerRS** | 6.7 | 5 | | | |
| **ThrRS** | 6.7 | 0 | | | |
| **TrpRS** | 6.7 | 0 | | | |
| **TyrRS** | 6.7 | 0 | | | |
| **ValRS** | 6.7 | 5 | | | |

**Table S2. Gene and seed concentration for all translation proteins.** Gene concentrations for a DNA brush with all translation genes. The solution is diluted to the desired concentration of genes in the DNA brush with 1xPBS buffer, 7% glycerol, and a control gene coding for a non-interacting protein so that the total DNA concentration in the solution remains 150 nM. The second column provides the concentration of the protein seeds used in the PURE system for the DNA brush experiments in Fig. 4.

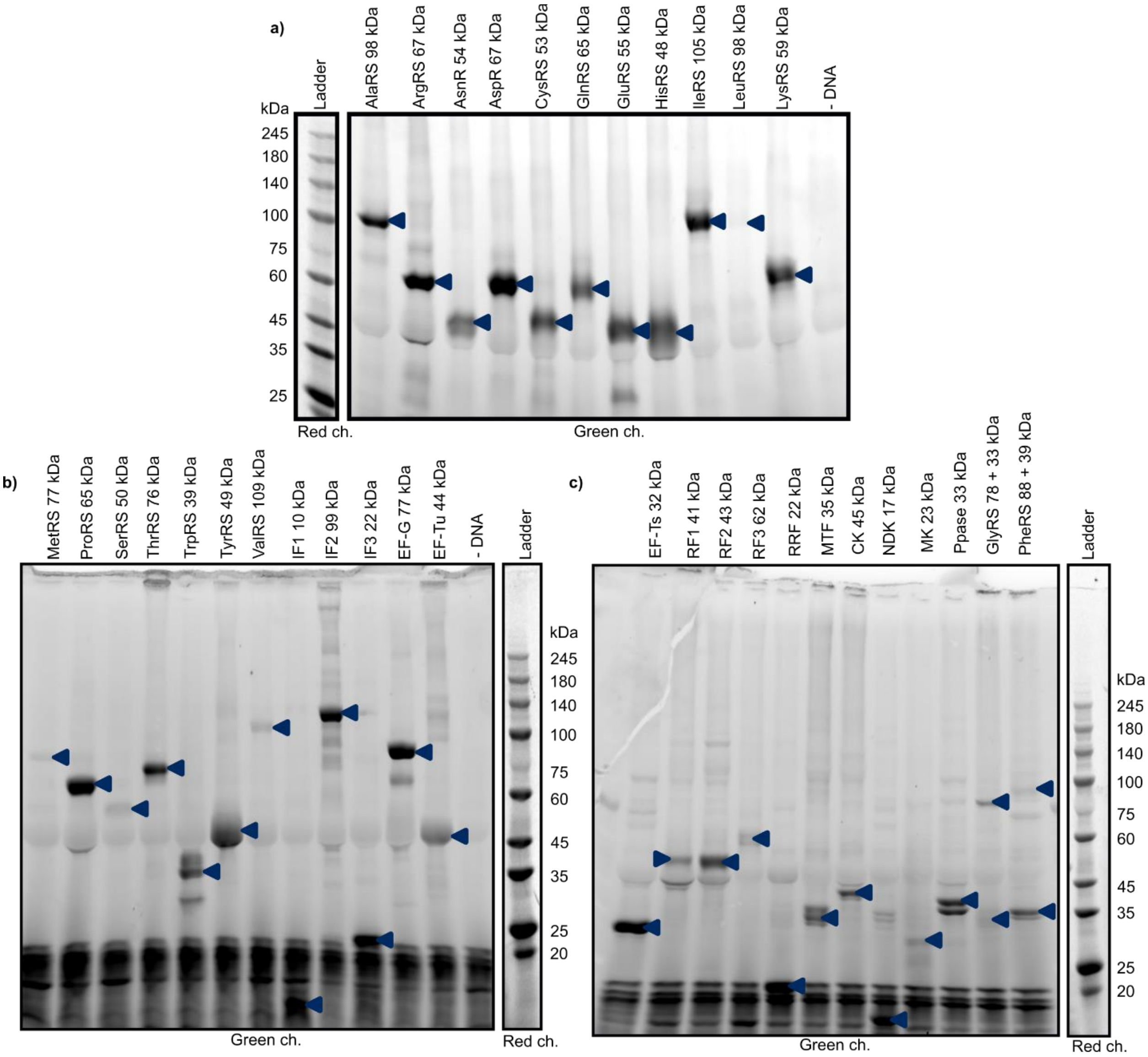

**Figure S1. Fluorescently labeled nascent PURE proteins resolved by SDS-PAGE. a)-c)** PURE reactions expressing each of the 36 PURE proteins were resolved by 4%-20% gradient SDS-PAGE (Methods). Green-Lys charged tRNA reagent was added to a 5 µl PURE reaction with the 1 nM of DNA coding for the corresponding PURE protein (4h, 37°C). After heat treatment (2min at 60°C) and SDS-PAGE, gels were imaged with a gel scanner (Methods). Fluorescent bands appearing at the expected molecular weight of expressed PURE protein are marked with arrows.

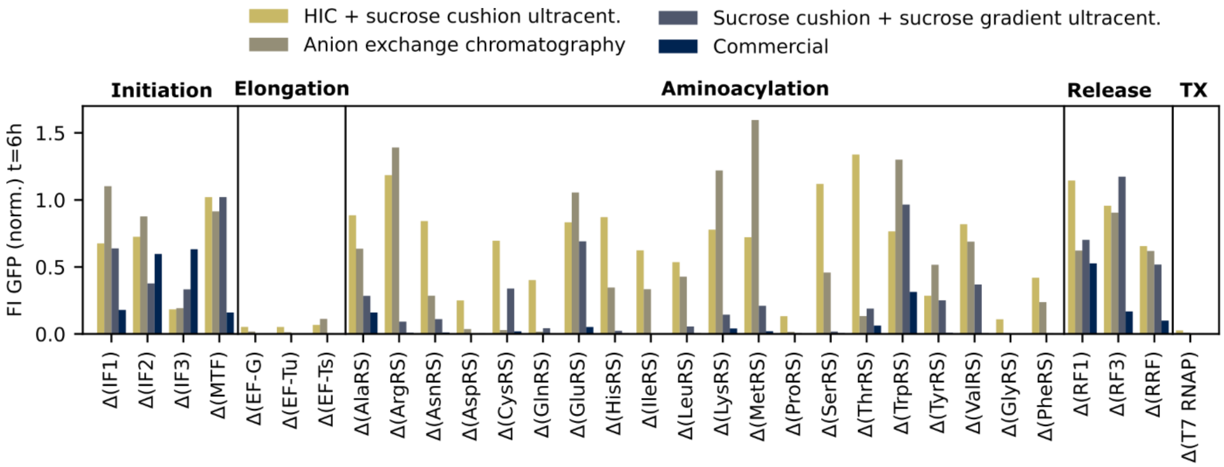

**Figure S2. Ribosome purity test.** GFP Fluorescent intensities (FI) at t=6h for different delta versions of the PURE with purified ribosomes from different methods. Method 1: Hydrophobic interaction chromatography (HIC) with subsequent sucrose cushion ultracentrifugation step [1]. Method 2: One-step anion exchange chromatography [2]. Method 3: Sucrose cushion ultracentrifugation with following sucrose gradient ultracentrifugation step with isolation of a peak at λ = 260 nm [3]. Method 4: Commercial ribosomes from PUREfrex 2.0 (Cosmo Bio USA) without DnaK/J and GroEL/S mix. GFP FI. are normalized to the FI of a full PURE sample in the same experiment. For more details, see Material and Methods.

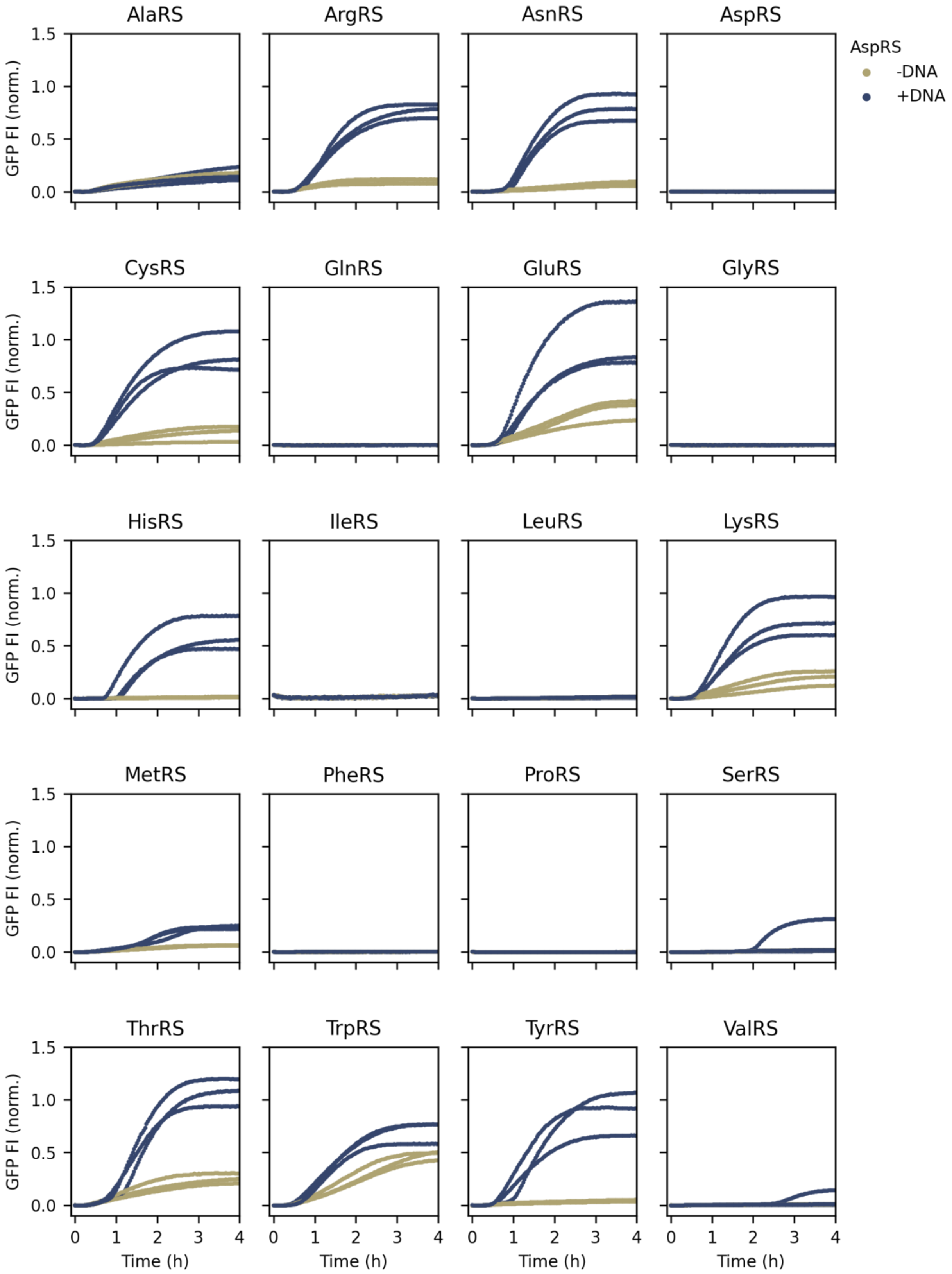
AlaRS
ArgRS
AsnRS
AspRS
AspRS
-DNA
+DNA
CysRS
GlnRS
GluRS
GlyRS
HisRS
IleRS
LeuRS
LysRS
MetRS
PheRS
ProRS
SerRS
ThrRS
TrpRS
TyrRS
ValRS
GFP FI (norm.)
1.5
1.0
0.5
0.0
0
1
2
3
4
Time (h)

**Figure S3. AaRS-based recovery in ΔPi-PURE.** Three repeats of Fl time traces, FI(t), of $\Delta P_i$-PURE for each of the 20 aaRS with and without the gene coding for the aaRS. No external seed was added. FI are normalized to the max. int. of a full PURE sample. For DNA concentrations, see Table S1.

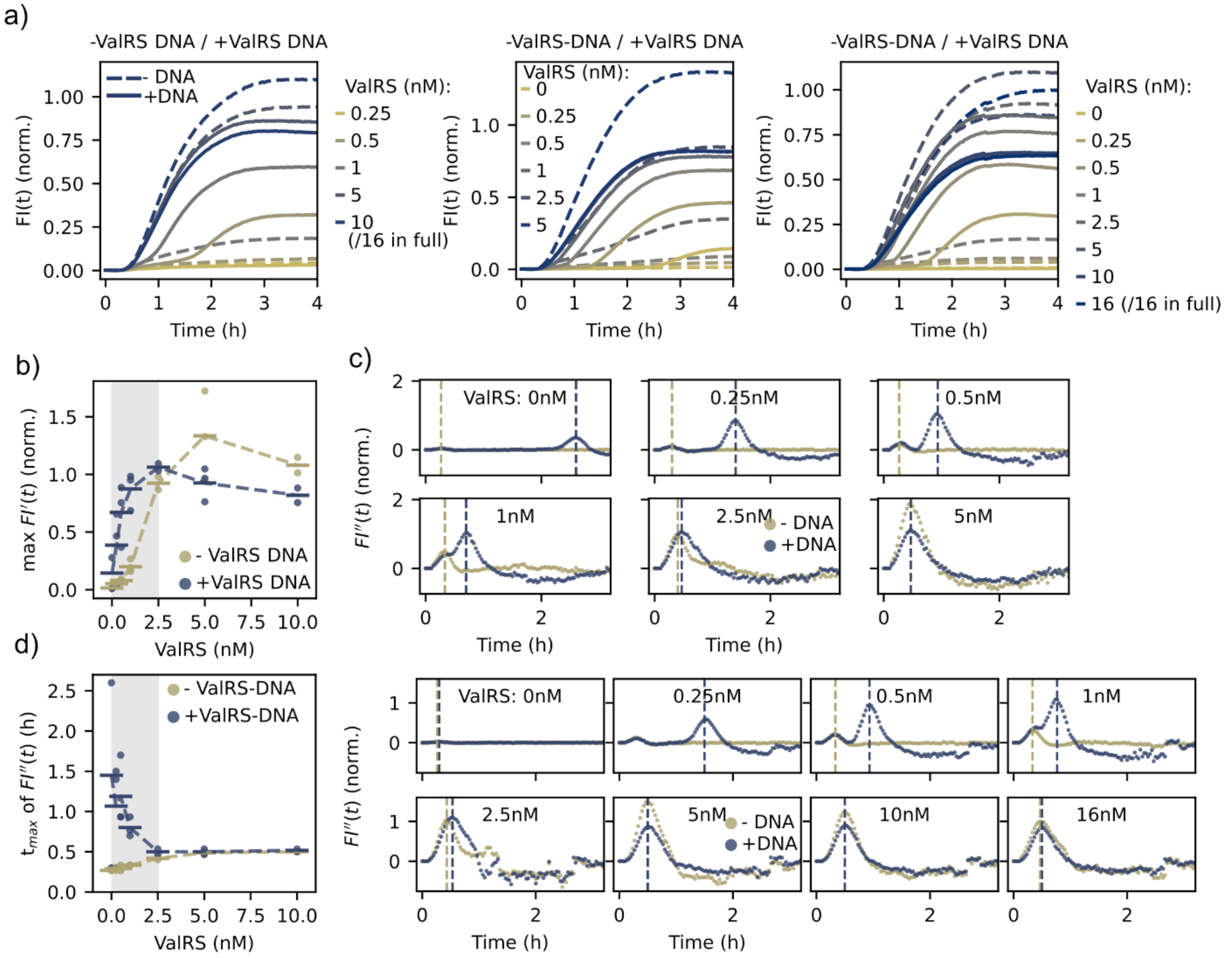

**Figure S4. Seeded recovery with ValRS.** a) Three repeats of Fl time traces, FI(t), of ΔValRS-PURE with different amounts of purified ValRS. Fl. Int. are normalized to the max. int. of a full PURE sample. $[DNA_{GFP}]$ = 1 nM, $[DNA_{ValRS}]$ = 1.5 nM. b) Max. of the first derivative of FI(t) as a function of the ValRS seed. Max. rate is normalized to the max. rate of a full PURE. Mean values are shown as horizontal markers and are connected via a dotted line. n=3. c) Second derivative of FI(t) of the two repeats with respect to time for different seed concentrations +/- DNA that is not shown in Fig. 1d. Dashed lines at time points of max. values of FI´´(t) as a guide to the eye. Values are normalized to the max. FI´´(t) of a full PURE sample. d) tmax is the time of max. FI´´(t) as a function of ValRS. n=3.

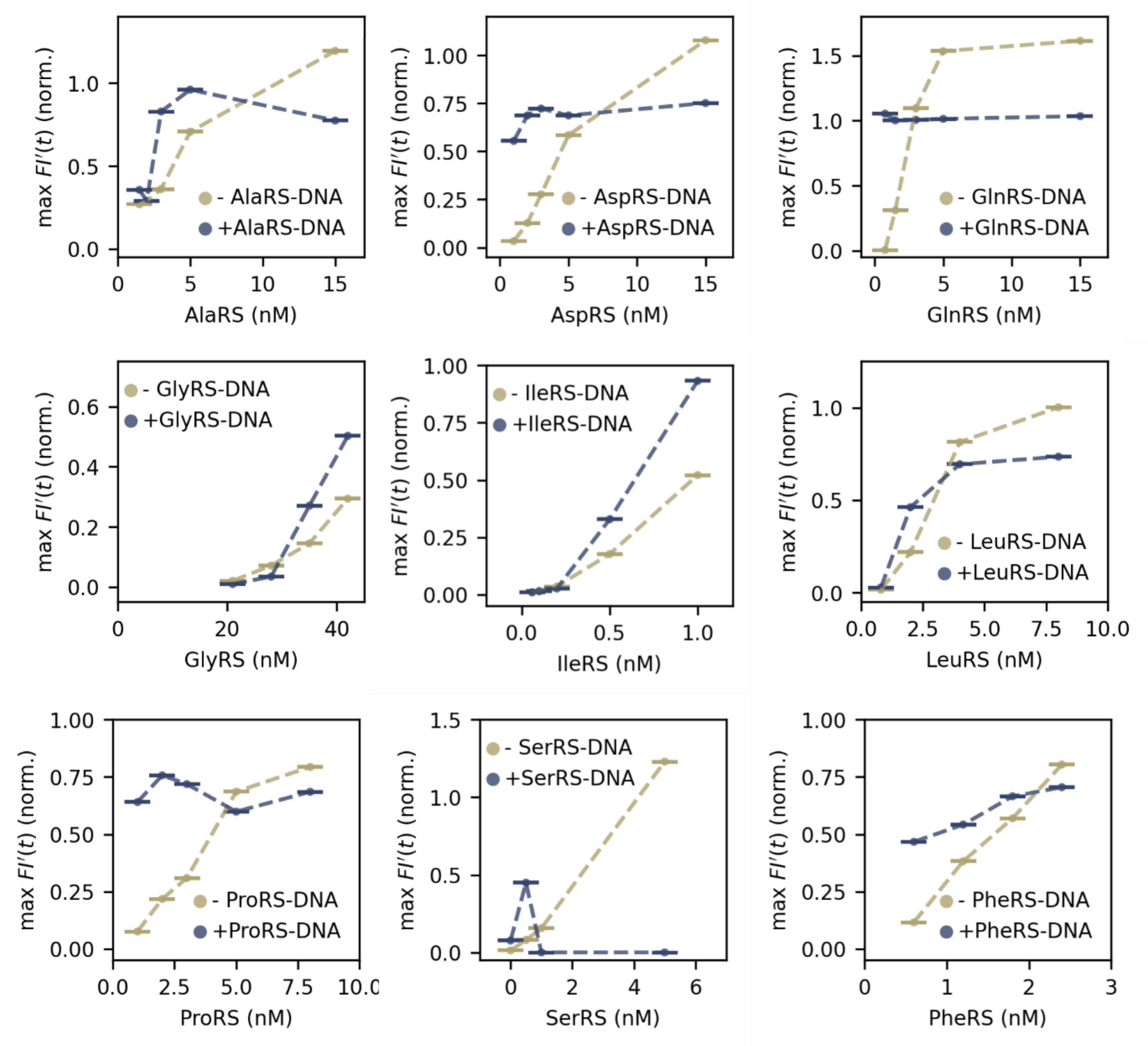

**Figure S5. Initial titration of aaRS for recovery assays.** Max. rate of GFP FI(t), from $\Delta P_i$-PURE with different amounts of purified aaRS with and without $P_i$-DNA. Int. are normalized to the max. rate of a full PURE sample. Based on these screens, we identified seed concentrations for Fig. 1e and S6.

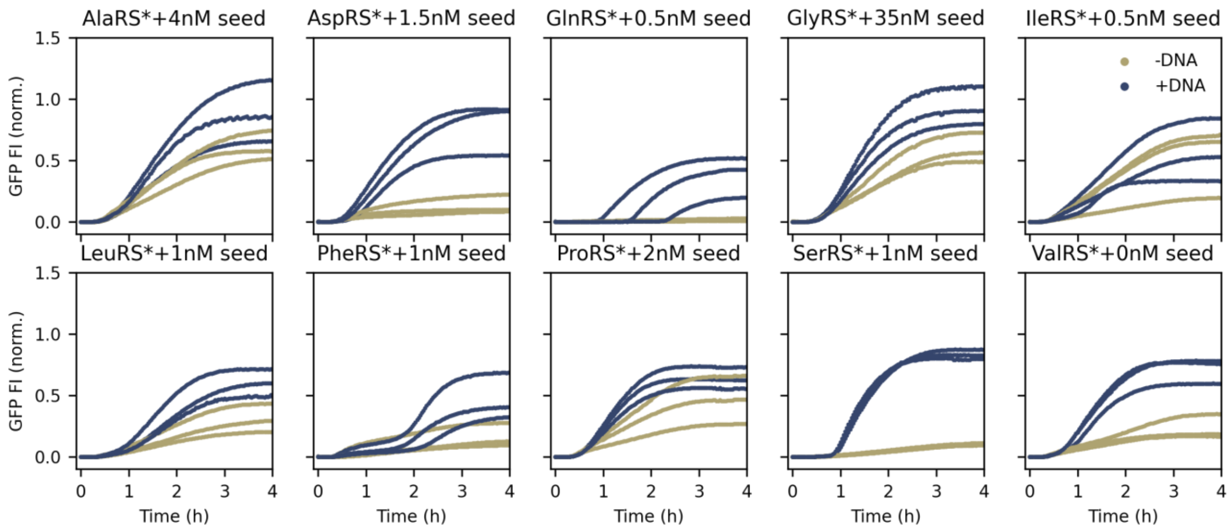

**Figure S6. Seeded recovery assay for 10 aaRS.** Three repeats of fluorescence intensity time traces, FI(t), of $\Delta P_i$-PURE with a seed for the 10 aaRS with and without the gene coding for the aaRS. Fl are normalized to the max. Fl of a full PURE sample. For DNA concentrations see Table S1.

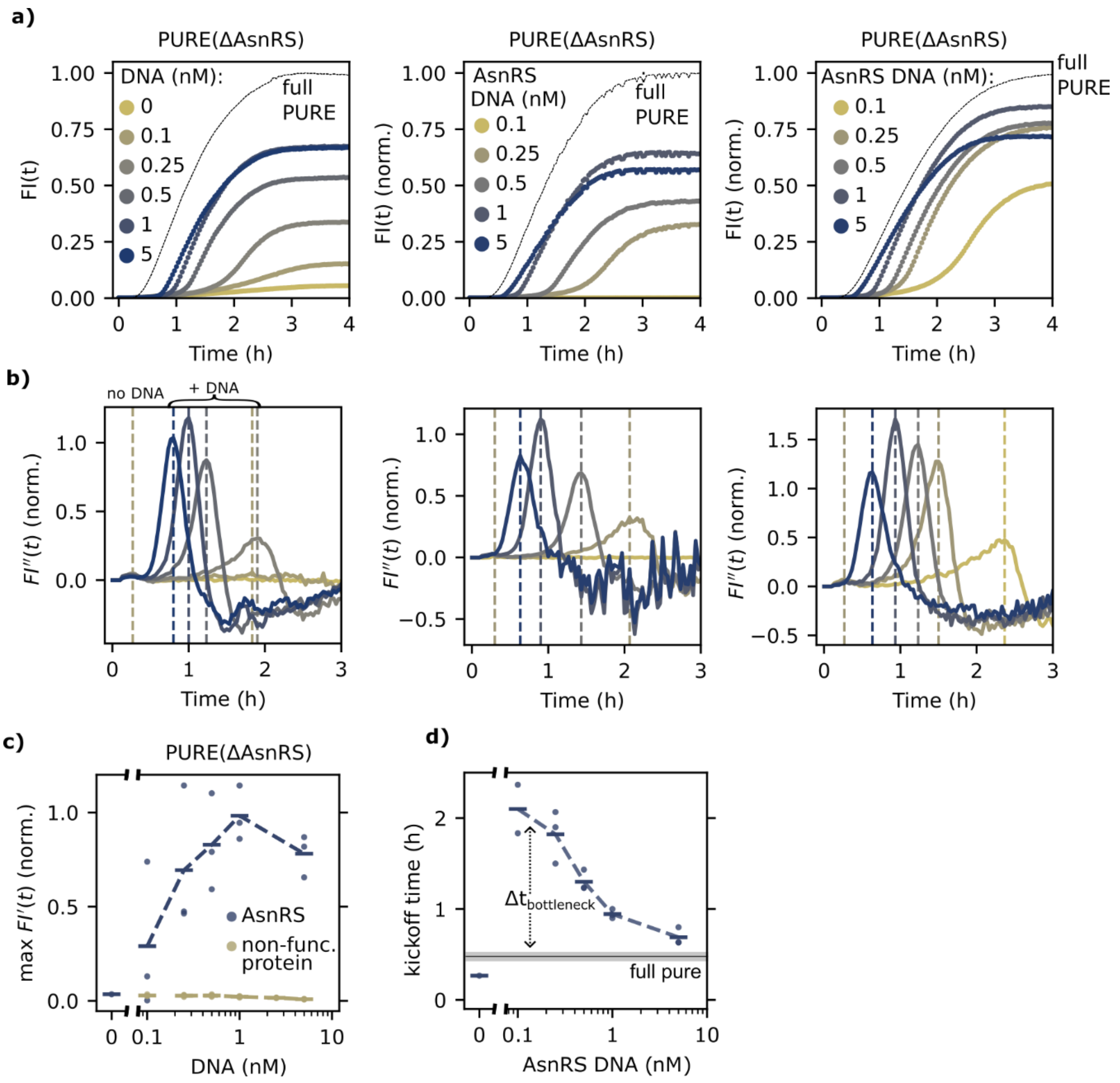

**Figure S7. Recovery assay for AsnRS.** a) Three repeats of FI(t) of ΔAsnRS-PURE with different amounts of DNA coding for AsnRS. FI are normalized to the max. FI of a full PURE sample (small black dots). [$DNA_{GFP}$] = 1 nM. b) Second derivative of FI(t) with respect to time for different [$DNA_{AsnRS}$]. Dashed lines at time points of max. values of FI´´(t) as a guide to the eye. Values are normalized to the max. FI´´(t) of a full PURE sample. The color code corresponds to the FI(t) curves above. C) Max. of the first derivative of FI(t) as a function of the DNA conc. for AsnRS. Max. rate is normalized to the max. rate of a full PURE sample. Mean values are shown as horizontal markers and are connected via a dotted line. d) The kickoff time is the time of max. FI´´(t) as a function of [$DNA_{AsnRS}$]. Max. FI´´(t) for a full PURE reaction shown as a solid line: mean. Grey area: std, n=3, fewer data points mean the absence of peaks above background noise.

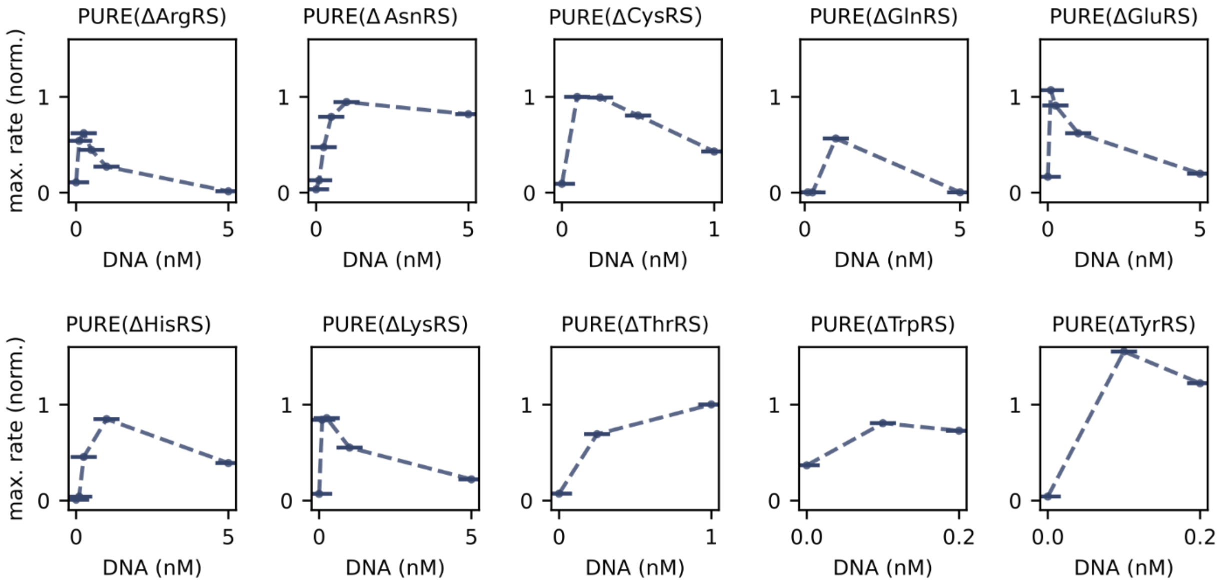

**Figure S8. Initial titrations of the aaRS genes for recovery.** Initial screening for DNA concentrations. $\Delta aaRS_i$-PURE with different amounts of DNA coding for $aaSR_i$ that did not require an external seed for recovery. Max. of the first derivative of fluorescence time traces as a function of the DNA conc. Max. rate is normalized to the max. rate of a full PURE sample.

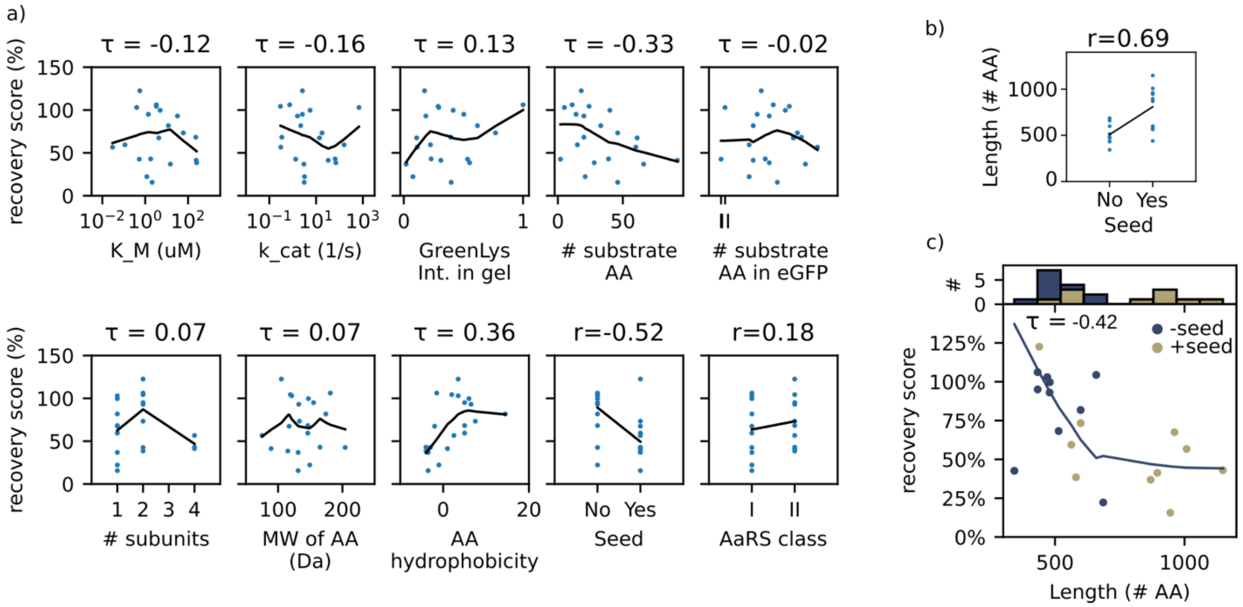

**Figure S9. Correlations with the recovery score. a)** The average recovery scores for all twenty aaRSs. were plotted as a function of several biochemical properties: The substrate affinity $K_M$, the enzyme turnover $k_{cat}$, size of one subunits [4], the number of amino acids $(AA)_i$ in the $aaRS_i$, requirement of a seed for PURE recovery, the oligomeric structure (1 for monomer, 2 for dimer, etc.) [5], the molecular weight of the AA, and the hydrophobicity score of the AA (the higher the score, the less hydrophobic) [6]. For each subplot, either Kendall's rank correlation coefficient $\tau$ or rank-biserial correlation *r* (for nominal data) is provided on top of each plot. **c)** Rank-biserial correlation r shown for the length of the aaRS (in #AA) dependent on the requirement of an external seed. **c)** The scores are shown for all twenty aaRSs as a function of their gene length (ORF). The solid lines fit the data (LOESS curves) and serve as guides to the eye.

**a)**

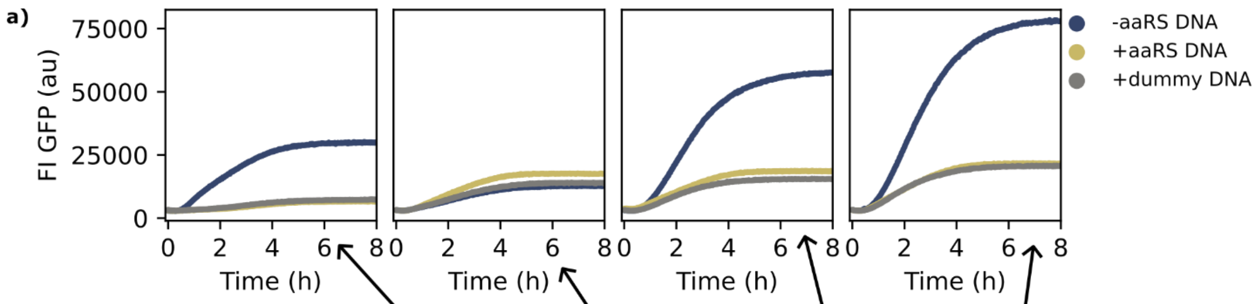

| Name | DNA (nM) | Seed (nM) | | | |
|---|---|---|---|---|---|
| | | Reaction 1 | Reaction 2 | Reaction 3 | Reaction 4 |
| AlaRS | 0.1 | 3.0 | 3.0 | 3.0 | 3.0 |
| ArgRS | 0.1 | 0.0 | 0.5 | 1.0 | 2.0 |
| AsnRS | 0.1 | 0.5 | 0.5 | 1.0 | 2.0 |
| AspRS | 0.1 | 1.5 | 1.5 | 1.5 | 1.5 |
| CysRS | 0.1 | 0.0 | 0.5 | 1.0 | 2.0 |
| GlnRS | 0.1 | 0.8 | 0.8 | 1.0 | 2.0 |
| GluRS | 0.1 | 0.0 | 0.5 | 1.0 | 2.0 |
| GlyRS | 0.1 | 35.0 | 35.0 | 35.0 | 35.0 |
| HisRS | 0.1 | 0.5 | 0.5 | 1.0 | 2.0 |
| IleRS | 0.1 | 0.4 | 0.5 | 1.0 | 2.0 |
| LeuRS | 0.1 | 1.5 | 1.5 | 1.5 | 1.5 |
| LysRS | 0.1 | 0.0 | 0.5 | 1.0 | 2.0 |
| MetRS | 0.5 | 1.0 | 1.0 | 1.0 | 2.0 |
| PheRS | 0.1 | 1.0 | 1.0 | 1.0 | 2.0 |
| ProRS | 0.1 | 1.0 | 1.0 | 1.0 | 2.0 |
| SerRS | 0.1 | 1.0 | 1.0 | 1.0 | 2.0 |
| ThrRS | 0.1 | 0.0 | 0.5 | 1.0 | 2.0 |
| TrpRS | 0.1 | 0.0 | 0.5 | 1.0 | 2.0 |
| TyrRS | 0.1 | 0.0 | 0.5 | 1.0 | 2.0 |
| ValRS | 0.1 | 1.0 | 1.0 | 1.0 | 2.0 |
| dummy gene (gp8) | 2.4 | | | | |
| eGFP DNA | 0.1 | | | | |

**b)**

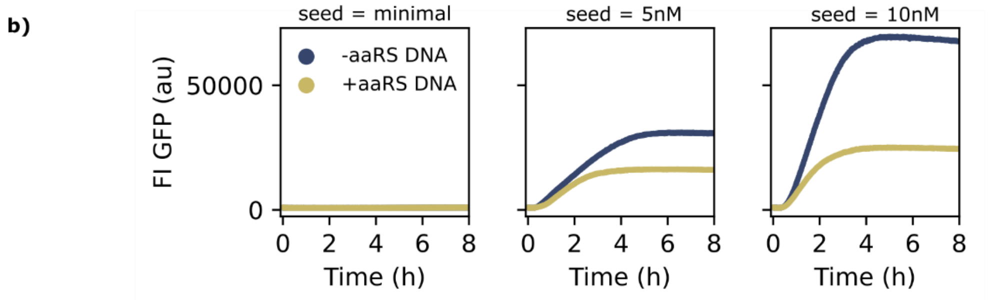

**Figure S10. Seeded recovery of twenty aaRSs in bulk solution reaction.** Measurements were performed in microplate wells. a) FI(t) for different ΔaaRS-PURE with different amounts of purified aaRS's and aaRS genes (concentrations are provided in the table). The GFP signal is lower in the presence of either aaRS DNA, or a non-interacting control gene, than without any additional DNA besides GFP DNA. b) Recovery for eighteen aaRS (w/o GlyRS, w/o PheRS), for three different seed concentrations with and without corresponding aaRS genes. Minimum seed (nM): AlaRS = 3.0, AspRS = 2.0, GlnRS = 0.8, IleRs = 0.5, LeuRS = 2.0, MetRs = 0.5, ProRs = 1, SerRS = 1, ValRs = 1. [aaRS-DNA] = 0.1 nM (except for MetRS-DNA = 0.4 nM), [GFP-DNA] = 0.4 nM. Other seeds 5nM or 10 nM for all aaRS.

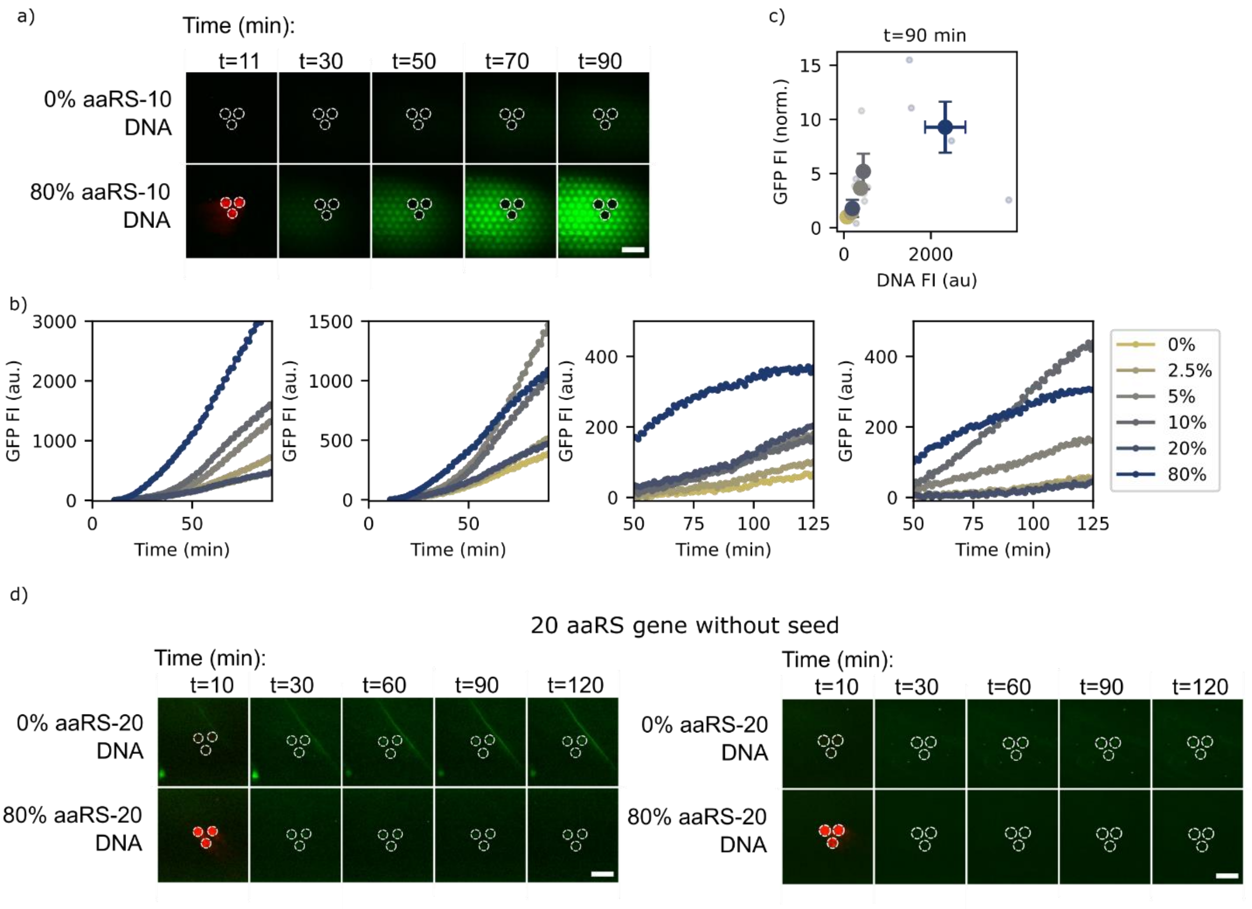

**Figure S11. Initial recovery assay of aaRSs groups from DNA brushes**. a-c) Biogenesis of ten aaRS (for Arg, Asn, Cys, Glu, His, Lys, Met, Ser, Thr, Trp, Tyr) from ΔaaRS$_{1\text{-}10}$-PURE frp, a DNA brush without seed. a) TIRF microscopy images of a hexagon-patterned surface traps for GFP reporter capturing (green channel) and three-DNA brush clusters (white dotted circles). All DNA brushes contain 2% GFP-DNA (labeled) and 80% genes coding to a non-interacting control protein (top) or ten aaRSs. Scale bar = 200 µm, b) Average GFP Fl(t) from TIRF microscopy images for DNA brushes with different fractions of the ten aaRS genes. c) Normalized average GFP Fl at t=90 min from TIRF microscopy as a function of the DNA Fl (aaRS DNA + GFP DNA) for different DNA brushes. n=4. Values are normalized to the average GFP Fl of a DNA brush with 0% aaRS genes. d) Attempt for recovery of twenty aaRSs from a DNA brush without any seed shows no GFP Fl with time. Two repeats of the same experiment. Image labels as in (a). All DNA brushes contain 2% GFP-DNA (labeled). We did not detect any accumulation of surface-bound GFP-HA as a function of the fraction of aaRS genes in the DNA brush. Scale bar = 200 um.

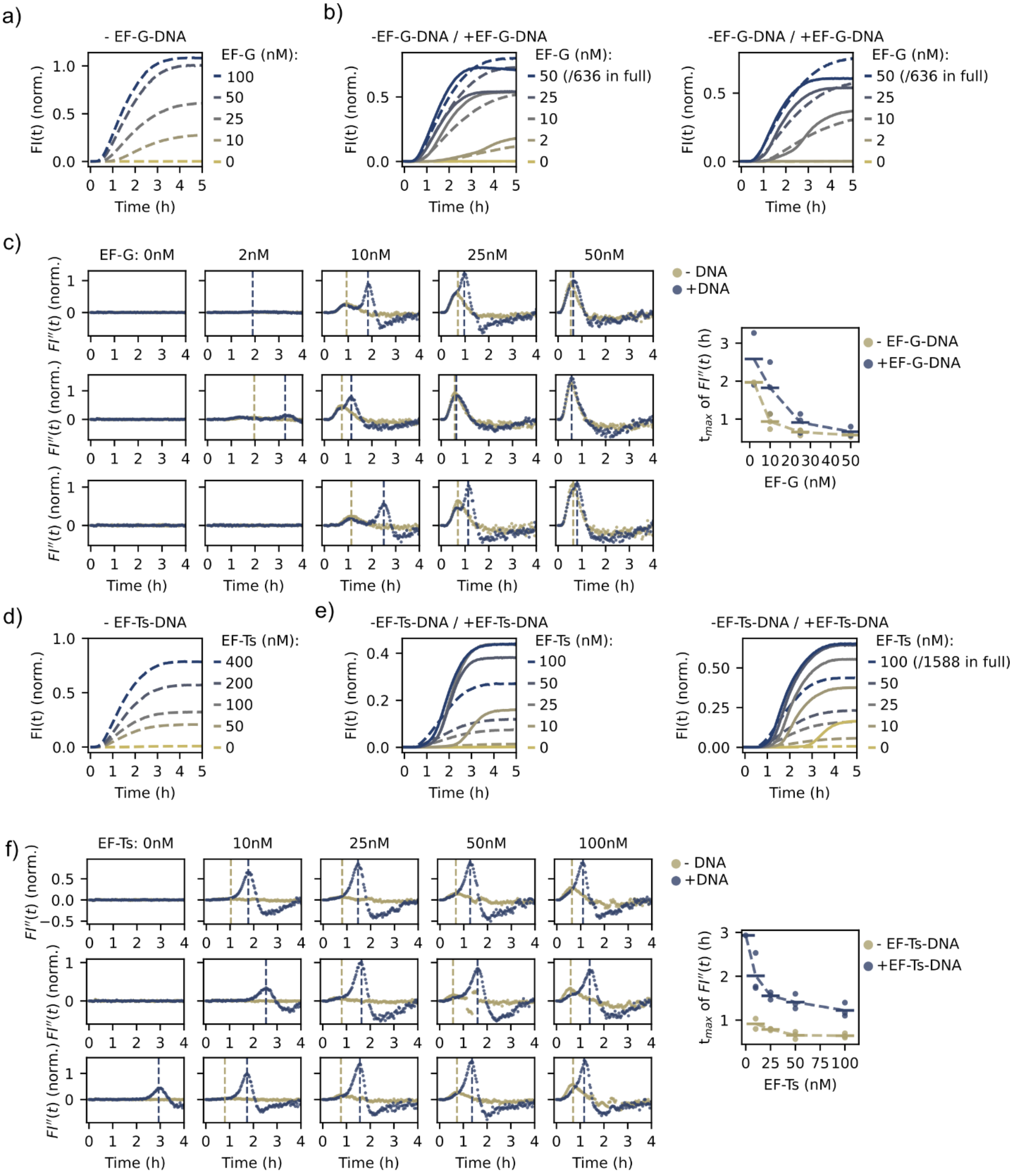

**Figure S12. Recovery tests for EF-G and EF-Ts in bulk solution.** Measurements were performed in a 384-well microplate. a) Fl(t) of GFP expression for an initial titration of EF-G in a ΔEF-G-PURE b) Two more repeats of the fluorescence intensity time traces of ΔEF-G-PURE with different amounts of EF-G. Fl are normalized to the max. Fl of a full PURE sample. $[DNA_{GFP}]$ = 1 nM, $[DNA_{EF\text{-}G}]$ = 0.75 nM. Dashed lines: without DNA, solid lines: with DNA. c) The second derivative of FI(t) with respect to time for different seed concentrations +/- DNA of the three repeats. Dashed lines at time points of max. values of F´´(t) as a guide to the eye. Values are normalized to the max. F´´(t) of a full PURE sample. Right: $t_{max}$ is the time of max. F´´(t) as a function of EF-G. n=3, less data points mean the absence of peaks above background noise. d) Fl(t) of GFP expression for an initial titration of EF-Ts in a ΔEF-Ts-PURE e) Two more repeats of the fluorescence intensity time traces of ΔEF-Ts-PURE with different amounts of EF-Ts. Fl are normalized to the max. Fl of a full PURE sample. [$DNA_{GFP}$] = 1 nM, [$DNA_{EF\text{-}Ts}$] = 1 nM. f) The second derivative of FI(t) with respect to time for different seed concentrations +/- DNA of the three repeats. Dashed lines at time points of max. values of F´´(t) as a guide to the eye. Values are normalized to the max. F´´(t) of a full PURE sample. Right: $t_{max}$ is the time of max. F´´(t) as a function of EF-Ts. n=3, less data points mean the absence of peaks above background noise.

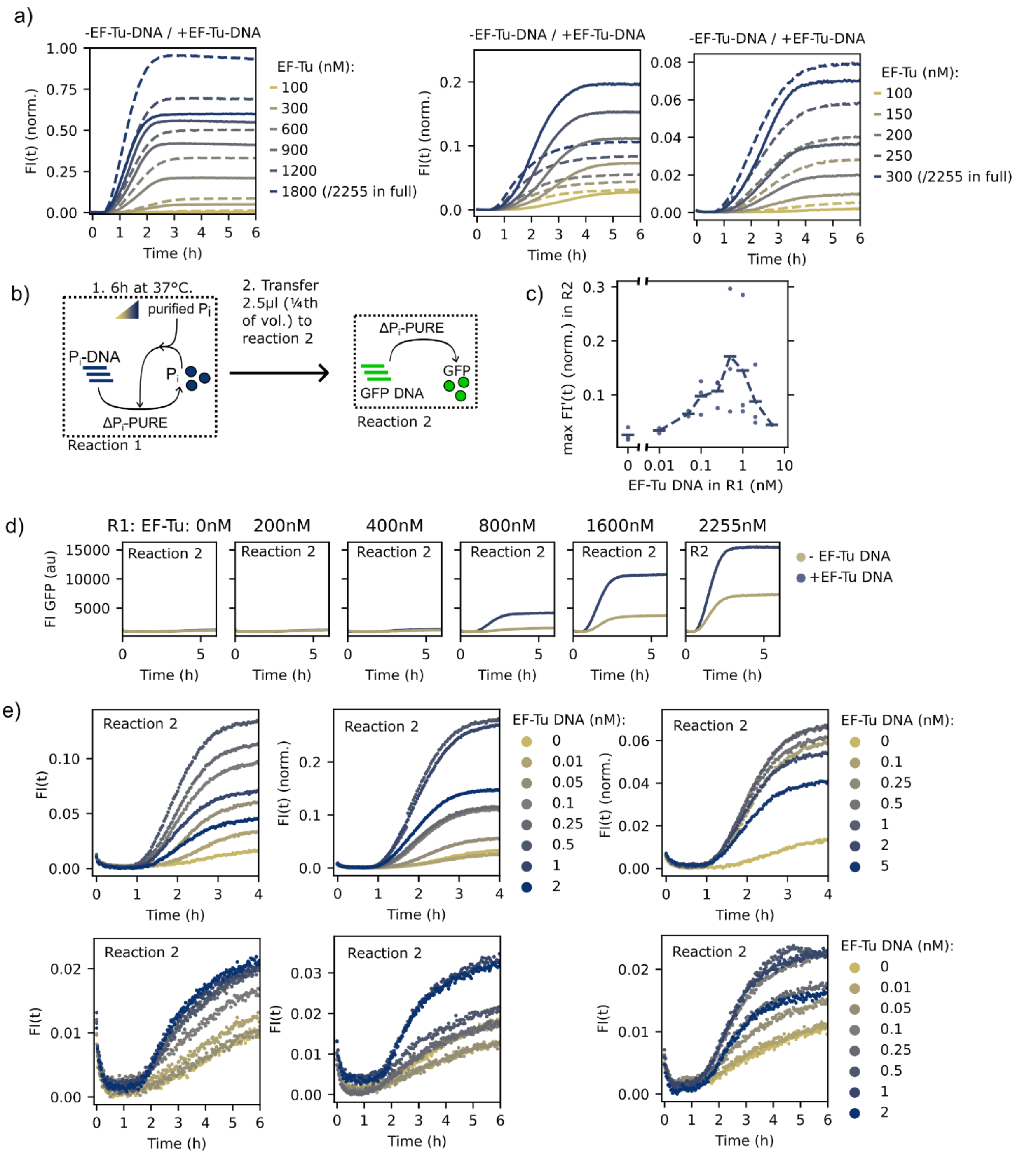

**Figure S13. EF-Tu recovery test in bulk solution.** Measurements were performed in a 384-well microplate. a) Three repeats of the Fl(t) of ΔEF-Tu-PURE with different amounts of EF-Tu. Fl are normalized to the max. Fl of a full PURE sample. $[DNA_{GFP}]$ = 1 nM, $[DNA_{EF\text{-}Tu}]$ = 1.5 nM. Dashed lines: without EF-Tu DNA, solid lines: with EF-Tu DNA. Left: GFP signal with EF-Tu DNA is higher than without DNA. Middle and left: GFP signal is lower with EF-Tu DNA than without DNA. b) Scheme of the recovery assay in two steps. Reaction 1 (R1): expression of EF-Tu from DNA with differently seeded PURE. Reaction 2: transfer from R1 into ΔEF-Tu-PURE with GFP DNA. c) Max. of the first derivative of Fl(t) from R2 as a function of the EF-Tu

DNA in R1. Max. rate is normalized to the max. rate of a full PURE sample. Mean values (horizontal markers) for n=6. d) Fl(t) of R2 spiked with 2.5 ul of R1 containing different amounts of EF-Tu. Normalized to the max. Fl of a full PURE sample. [$DNA_{EF\text{-}Tu}$] = [$DNA_{GFP}$] = 0.4 nM. d) Fl(t) from two repeats for reaction 2 spiked with 2.5 ul of R1 containing different amounts of EF-Tu DNA. Normalized to the max. Fl of a full PURE sample. [$DNA_{GFP}$] = 0.4 nM, seed = 800 nM EF-Tu. e) Fl(t) from three repeats of R2 spiked with 2.5 ul of R1 containing different amounts of EF-Tu DNA. Normalized to the max. Fl of a full PURE sample. [$DNA_{GFP}$] = 0.4 nM, seed = 800 nM EF-Tu.

a)

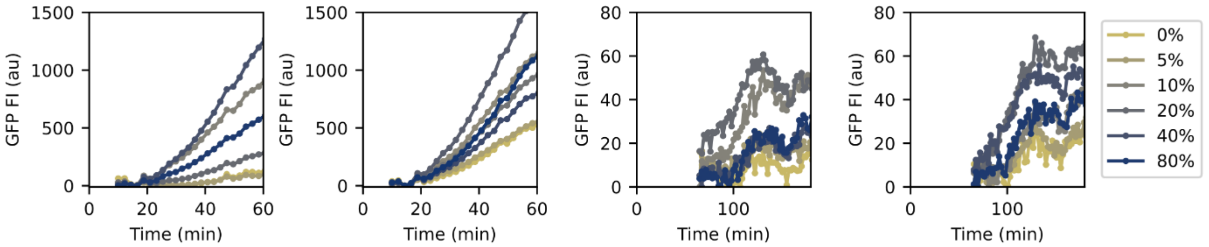

b)

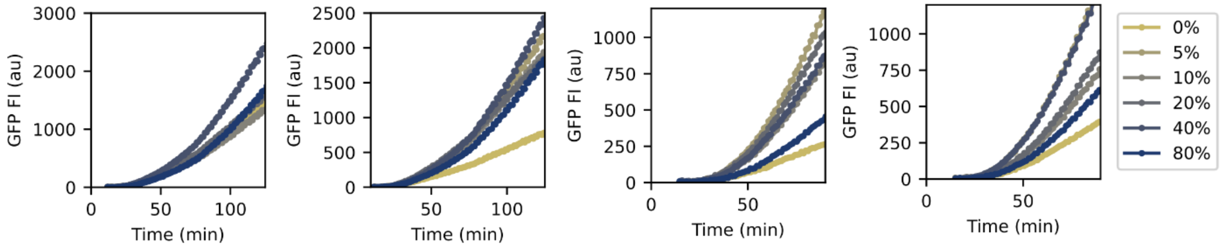

**Figure S14. EF-Tu and EFs biogenesis from DNA brushes.** a) Fl(t) of average GFP fluorescence from TIRF microscopy images for DNA brushes with different fractions of EF-TU gene for data shown in Figure 1g. b) Fl(t) of average GFP fluorescence from TIRF microscopy images for DNA brushes with different fractions of EF group genes for data shown in Figure 1j.

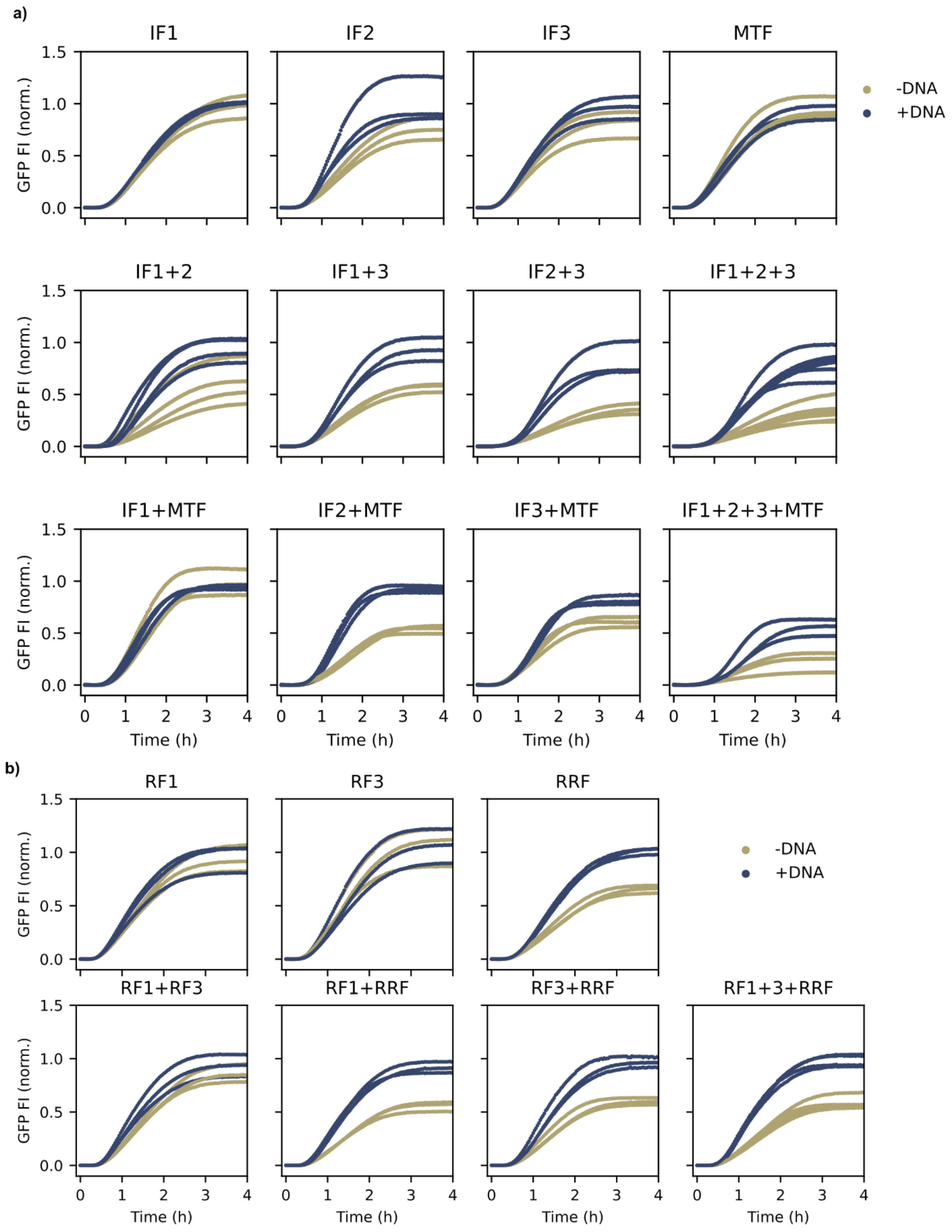

**Figure S15. Recovery of the initiation and release factors in bulk solution.** a) Minimum three repeats of FI(t) of ΔIF$_i$-PURE for all the permutations of double deletions with the three IFs and methionyl-tRNA formyltransferase (MTF) +/- their corresponding DNA. For each IF: [$DNA_{IF}$] = 0.1 nM and [$DNA_{GFP}$] = 0.4 nM, Fl are normalized to the Fl of a full PURE sample. **b)** Replicates for the recovery test for the release and recycling factors without a seed. Repeats of FI(t) for all the grouping of RF1, RF3 and RRF -/+ their corresponding DNA. For each RF: [$DNA_{RF}$] = 0.1 nM and [$DNA_{GFP}$] = 0.4 nM, minimum n=3. Fl are normalized to the max. Fl of a full PURE sample. Measurements were performed in a 384-well microplate.

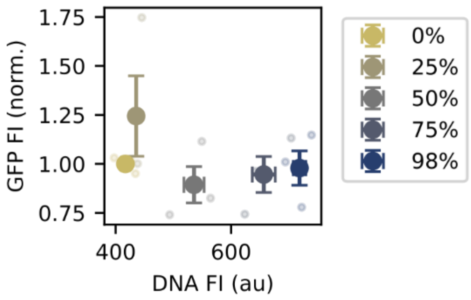

**Figure S16. DNA brush with all translation genes with insufficient seed.** a) Normalized mean GFP Fl and its s.e.m. from TIRF microscopy images as a function of the DNA fluorescence signal (translation protein-DNA + GFP DNA) for different DNA brushes. n=3. Values are normalized to the average GFP Fl of a DNA brush with 0% translation genes, 2% GFP-HA genes, and 98% dummy gene. PURE system contains EF-G = 25 nM, EF-Tu=1000 nM, EF-Ts=100 nM, each aaRS = 10 nM, except for GlyRS = 40 nM and PheRS = 20 nM. No seed for IF nor RF. No significant correlation between the composition of the genes in the DNA brush and the GFP Fl was observed. See Table S2 for gene concentrations in the DNA brush.

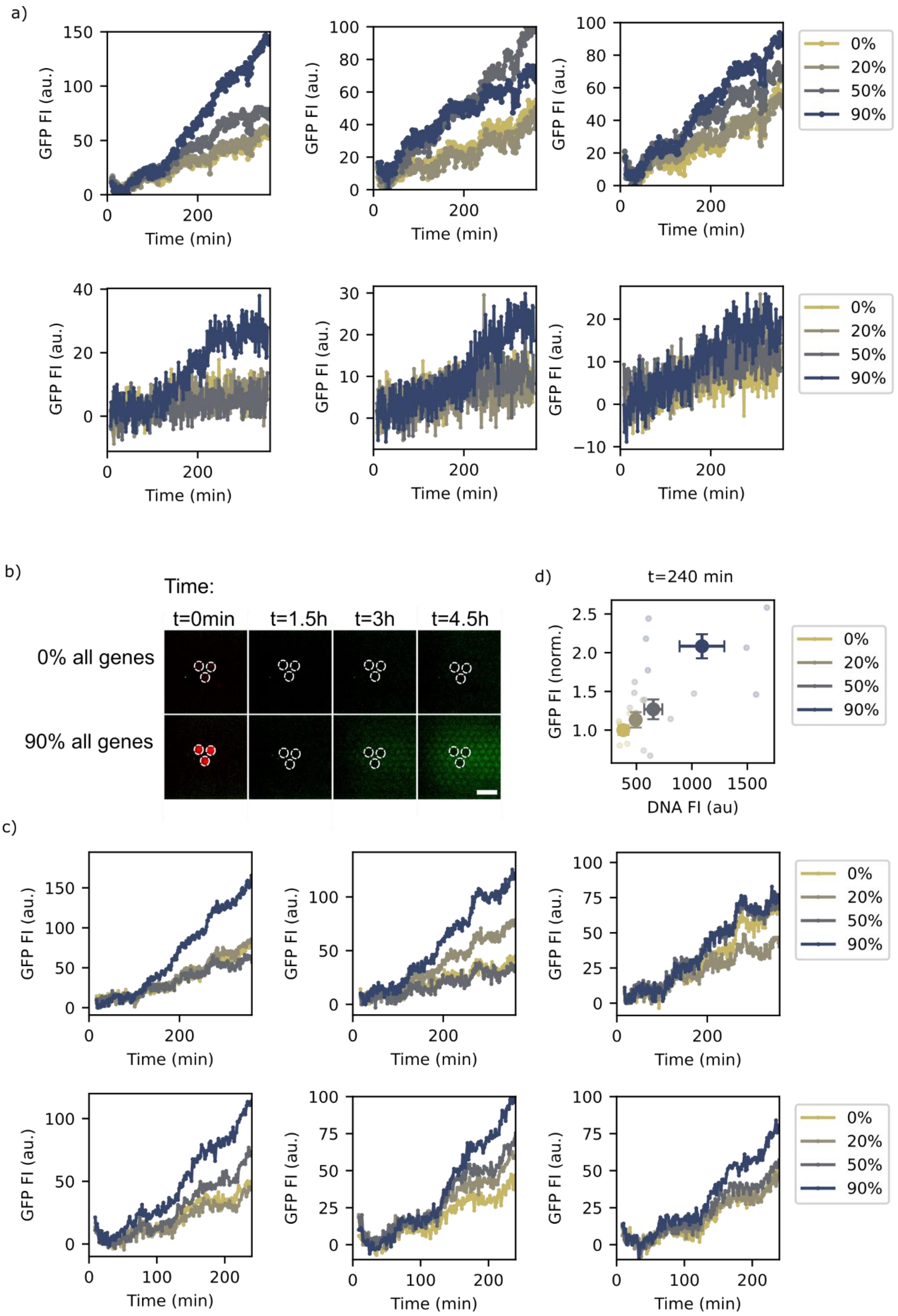
a)
GFP FI (au.)
Time (min)
0%
20%
50%
90%
b)
Time:
t=0min
t=1.5h
t=3h
t=4.5h
0% all genes
90% all genes
c)
d)
t=240 min
GFP FI (norm.)
DNA FI (au)

**Figure S17. Biogenesis of all translation proteins from DNA brushes with different seeds.** a) Average GFP Fl(t) from TIRF microscopy images for DNA brushes. The PURE system contains RF seed, IF seed, EF seed, and ten aaRS seed (see Table S2 and Fig. 4. b-d) Biogenesis of all thirty translation proteins from a DNA brush with a higher seed concentration. Seed for the aaRS, IF and RF are 2x the concentration listed in Table S2. Seed for EF-G = 50nM, EF-Ts = 200nM, EF-Tu = 1000nM. b) TIRF microscopy images of a hexagon-patterned surface traps for GFP reporter capturing (green channel) and three-DNA brush clusters (white dotted circles). The bottom left image shows the fluorescently labeled DNA (red channel). The DNA brushes contained 2% GFP genes and a total of 90 % genes either coding for a non-interacting control protein (DNA-not labeled, top row) or coding for all translation proteins (DNA-labeled, bottom row). The composition of the PURE system as in (a). Scale bar = 200 µm. c): Average GFP Fl(t) from TIRF microscopy images for DNA brushes with different fractions of translation genes. c) Normalized mean GFP Fl and its s.e.m. from TIRF microscopy as a function of the DNA Fl (translation protein-DNA + GFP DNA) for different DNA brushes. n=6. Values are normalized to the average GFP signal of a DNA brush with 0% translation genes.
